# Global diversity and biogeography of the *Zostera marina* mycobiome

**DOI:** 10.1101/2020.10.29.361022

**Authors:** Cassandra L. Ettinger, Laura E. Vann, Jonathan A. Eisen

## Abstract

Seagrasses are marine flowering plants that provide critical ecosystem services in coastal environments worldwide. Marine fungi are often overlooked in microbiome and seagrass studies, despite terrestrial fungi having critical functional roles as decomposers, pathogens or endophytes in global ecosystems. Here we characterize the distribution of fungi associated with the seagrass, *Zostera marina,* using leaves, roots, and rhizosphere sediment from 16 locations across its full biogeographic range. Using high throughput sequencing of the ribosomal internal transcribed spacer (ITS) region and 18S ribosomal RNA gene, we first measured fungal community composition and diversity, then we tested hypotheses of neutral community assembly theory and the degree to which deviations suggested amplicon sequence variants (ASVs) were plant-selected or dispersal-limited, and finally we identified a core mycobiome and investigated the global distribution of differentially abundant ASVs. Our results show that the fungal community is significantly different between sites and follows a weak, but significant pattern of distance decay. Generally, there was evidence for both deterministic and stochastic factors contributing to community assembly of the mycobiome. The *Z. marina* core leaf and root mycobiomes are dominated by unclassified Sordariomycetes spp., unclassified Chytridiomycota lineages (including Lobulomycetaceae spp.), unclassified Capnodiales spp. and *Saccharomyces* sp. A few ASVs (e.g. *Lobulomyces* sp.) appear restricted to one or a handful of locations (e.g. possibly due to local adaptation, deterministic dispersal limitation or seasonal bloom events), while others (e.g. *Saccharomyces* sp.) are more ubiquitous across all locations suggesting a true global distribution and possible plant-selection. Fungal guilds associated with *Z. marina* were only weakly identified (10.12% of ITS region and 3.4% 18S rRNA gene ASV guild assignments were considered highly probable) including wood saprotrophs, ectomycorrhizal fungi, endophytic fungi and plant pathogens. Our results are similar to those found for other seagrass species. It is clear from the many unclassified fungal ASVs and fungal functional guilds, that our knowledge of marine fungi is still rudimentary. Further studies characterizing seagrass-associated fungi are needed to understand the roles of these microorganisms generally and when associated with seagrasses.

## Introduction

Terrestrial fungi are known to have critical ecological roles as microbial saprotrophs, pathogens and mutualists (Peay *et al.*, 2016), and although less is known about fungi in aquatic ecosystems, it is believed that they also have vital ecological roles (e.g. in organic matter degradation, nutrient cycling and food web dynamics (Kagami *et al.*, 2007; Gutiérrez *et al.*, 2011; Orsi *et al.*, 2013; Grossart *et al.*, 2016, 2019; Raghukumar, 2017)). Despite their global importance, the taxonomic, phylogenetic, and functional diversity of marine fungi generally is vastly understudied (Amend *et al.*, 2019). In comparison to the greater than 120,000 terrestrial fungal species known (Hawksworth & Lücking, 2017), there are currently only ~1,692 described species of marine fungi, though estimates of the true diversity of these organisms is much higher (Jones, 2011; Jones *et al.*, 2015, 2019). Recent studies have examined the global distribution of marine planktonic, pelagic, and benthic fungi (Tisthammer *et al.*, 2016; Morales *et al.*, 2019; Hassett *et al.*, 2020), yet the distribution of host-associated fungi in the marine environment is still relatively unknown. Fungi have been reported in association with many marine animals including sponges (Gao *et al.*, 2008), corals (Littman *et al.*, 2011) and other invertebrates (Yarden, 2014), with algae and seaweeds (Zuccaro *et al.*, 2008; Gnavi *et al.*, 2017), and flowering plants, like seagrasses (Borovec & Vohník, 2018).

Seagrasses are foundation species in coastal ecosystems worldwide and are the only submerged angiosperms (flowering plants) to inhabit the marine environment. One widespread seagrass species, *Zostera marina*, also known as eelgrass, provides critical ecosystem services in coastal environments throughout much of the Northern Hemisphere (Hemminga & Duarte, 2000; Orth *et al.*, 2006; Fourqurean *et al.*, 2012). Previous studies have investigated the composition and structure of the bacterial community associated with *Z*. *marina*, including a global survey that was able to identify a core eelgrass root microbiome (Ettinger *et al.*, 2017a; Fahimipour *et al.*, 2017; Bengtsson *et al.*, 2017). Members of this community are thought to facilitate nitrogen and sulfur cycling for host plant benefit (Capone, 1982; Sun *et al.*, 2015; Cúcio *et al.*, 2016; Ettinger *et al.*, 2017a,b; Fahimipour *et al.*, 2017; Crump *et al.*, 2018; Wang *et al.*, 2020).

Comparatively, not as much is known about the distribution, diversity, and function of the mycobiome (i.e. the fungal community) associated with *Z*. *marina*. Culture-based studies have described a mycobiome composed of taxa in the classes Eurotiomycetes, Dothideomycetes, and Sordariomycetes (Shoemaker & Wyllie-Echeverria, 2013; Kirichuk & Pivkin, 2015; Petersen *et al.*, 2019; Ettinger & Eisen, 2020). These studies consistently find dominance of a few ubiquitous taxa (e.g. *Cladosporium* sp.) but also a diverse set of rare taxa that vary among sites and may be endemic to specific locations (e.g. *Colletotrichum* sp.) (Ettinger & Eisen, 2020). This pattern is suggestive of neutral community assembly through stochastic processes.

While culture-independent studies of *Z. marina* and other seagrass species have more exhaustively characterized the taxonomic diversity of these fungal communities, they have also highlighted how little is known about factors affecting the distribution, function and community assembly of seagrass-associated fungi (Wainwright *et al.*, 2018, 2019b; Hurtado-McCormick *et al.*, 2019; Ettinger & Eisen, 2019; Trevathan-Tackett *et al.*, 2020). A common finding among these studies is that taxonomic assignments cannot be made for greater than two-thirds of the fungal sequences associated with seagrasses and that Chytridiomycota lineages are dominant in this ecosystem (Wainwright *et al.*, 2019b; Ettinger & Eisen, 2019; Trevathan-Tackett *et al.*, 2020). Our culture-independent understanding of the mycobiome of *Z. marina* has so far focused on a single location in Bodega Bay, CA (Ettinger & Eisen, 2019). However, site-to-site variation in the mycobiome has now been observed in mycobiome studies from several other seagrass species (Wainwright *et al.*, 2018, 2019b; Hurtado-McCormick *et al.*, 2019; Trevathan-Tackett *et al.*, 2020) For example, a distance-decay relationship was found for the fungal community associated with the seagrass, *Enhalus acoroides*, in Singapore and Peninsular Malaysia (Wainwright *et al.*, 2019b), and for the seagrass, *Syringodium isoetifolium,* along Wallace’s line (Wainwright *et al.*, 2018). Additionally, the global planktonic marine fungal community was found to cluster by ocean (Hassett *et al.*, 2020), thus we might expect in our study, in addition to a distance-decay relationship, that we might see differentiation by ocean basin. Such geographic relationships are suggestive of niche-based community assembly through deterministic processes such as environmental filtering.

One concept central to our investigation here is the role of stochastic and deterministic drivers in determining the community assembly of the seagrass mycobiome. The Sloan neutral model has been widely applied to assess community assembly dynamics for microbial communities (Sloan *et al.*, 2007; Burns *et al.*, 2016). The assumption of this model is that random immigrations, births, and deaths can determine the relative abundance of taxa in a community (Sloan *et al.*, 2007). The model further assumes that local communities are assembled stochastically from regional pools, and that deterministic competitive interactions are not important in shaping the community because species are competitively equivalent (Chave, 2004; Rosindell *et al.*, 2011; Schmidt *et al.*, 2015). Stochastic processes supporting the neutral model include priority effects and ecological drift, while deterministic processes include species traits, interspecies interactions (e.g. competition, mutualisms) and environmental conditions (Zhou & Ning, 2017). Dispersal limitation can be either a stochastic or deterministic process (Lowe & McPeek, 2014). Identifying specific taxa that deviate from the model allows us to identify taxa that are assembled through deterministic processes including plant selection (Shade & Stopnisek, 2019).

Here we use high-throughput sequencing of marker genes to (1) characterize the fungal community associated with the seagrass, *Zostera marina,* globally and assess whether a distance-decay relationship is present between *Z. marina* and its mycobiome, (2) define a global core fungal community and assess community assembly dynamics using neutral models coupled with differential abundance analysis to predict important fungal taxa and evaluate their global distribution, and (3) assign functional predictions for the fungal community associated with *Z. marina*.

## Methods

### Sample collection

Samples were collected from 16 different globally distributed sites by researchers in the *Zostera* Experimental Network (ZEN) (Table S1) (Duffy *et al.*, 2015). Samples were collected subtidally at ~1 m depth using a modified version of the collection protocol previously used in Fahimipour et al. (2017). At each of the 16 sites, leaves and roots from individual *Z. marina* plants and adjacent sediment were collected for 12 individuals resulting in a total of 576 samples (n_leaf_ = 192, n_root_ = 192, n_sediment_ = 192).

To obtain *Z. marina* leaf and root tissues for analysis here, researchers were instructed to (1) gently remove individual *Z. marina* plants from the sediment, (2) briefly swish the individual in nearby seawater to remove loosely associated sediment from the roots, (3) collect ~5 roots and fully submerge in a pre-labelled 2 mL microcentrifuge tube filled with DNA/RNA Shield (Zymo Research, Inc, Irvine, CA, United States), and (4) collect a 2 cm section of healthy green leaf tissue and fully submerge in a pre-labelled 2 mL microcentrifuge tube filled with DNA/RNA Shield. A sample of sediment was taken adjacent to each *Z. marina* individual from 1 cm under the sediment surface using a 6CC syringe. Briefly this was performed by (1) removing the plunger from the syringe, (2) inserting the barrel of the syringe into the sediment, (3) inserting the syringe plunger to form an airtight seal, (4) removing the syringe from sediment, (5) extruding the sediment until the base of the syringe plunger is at the 3CC mark, and (6) using an alcohol sterilized plastic spatula to transfer ~0.25 g of sediment into a pre-labelled 2 mL microcentrifuge tube filled with DNA/RNA Shield. Samples were preserved in DNA/RNA Shield as it stabilizes DNA/RNA at room temperature. All samples were processed in the field immediately or within 5 hours of collection. Samples subsequently were kept at room temperature and mailed to the University of California, Davis within two weeks of sample collection.

### Molecular methods

Samples were shipped from UC Davis to Zymo Research, Inc. for DNA extraction. Samples were transferred to 96-well plate format, with plates including both positive (ZymoBIOMICS Microbial Community standard) and negative (no input) controls. DNA was extracted from samples using the ZymoBIOMICS DNA Miniprep kit following the manufacturer’s protocol with minor modifications as follows. Prior to DNA extraction, samples were heated at 65 °C for 5 minutes to resuspend any white precipitate that had accumulated. Sediment samples were vortexed for 30 seconds to ensure homogenization and then using a flame-sterilized spatula transferred into ZR BashingBead Lysis tubes until tubes were two-thirds full. Leaf and root samples were vortexed 30 seconds to dissociate any epiphytes and then all the liquid was transferred into ZR BashingBead Lysis tubes. For step 1, ZymoBIOMICS Lysis solution was then added to samples such that the final volume was ~1 mL. For step 2, samples were then subjected to a bead beater on the “homogenize” setting speed for 5 minutes. For step 4, 600 uL of supernatant was transferred to the filter tube. For step 11, only 50 uL of DNase/RNase free water was used for DNA elution. DNA concentrations for controls and a subset of samples per plate were first quantified with a Nanodrop (Thermo Fisher Scientific, Waltham, MA, United States), and subsequently all samples were quantified using Quant-iT PicoGreen (Thermo Fisher Scientific, Waltham, MA, United States). DNA was then shipped directly to the U.S. Department of Energy Joint Genome Institute (JGI) for amplicon sequencing.

### Sequence generation

The ribosomal internal transcribed spacer 2 (ITS2) region was amplified via polymerase chain reaction (PCR) using the ITS9F and ITS4R primer set (White *et al.*, 1990; Menkis *et al.*, 2012) and the 18S ribosomal RNA gene was amplified via PCR using the 565F and 948R primer set (Stoeck *et al.*, 2010). Libraries were prepared according to the JGI’s iTag library construction standard operating protocol (SOP) v.1.0 (https://1ofdmq2n8tc36m6i46scovo2e-wpengine.netdna-ssl.com/wp-content/uploads/2019/07/iTag-Sample-Preparation-for-Illumina-Sequencing-SOP-v1.0.pdf). We briefly summarize their protocol here. Three replicate PCR reactions for each sample were performed in 96-well plate format with the following conditions: 94 °C for 3 min, 35 cycles at 94 °C for 25 sec, 50 °C for 60 sec, 72 °C for 90 sec, and a final extension at 72 °C for 10 min. After amplification, replicate PCR products were combined and then samples were pooled together based on DNA quantification of combined PCR replicates. Samples were then pooled at up to 184 samples per sequencing run and sequenced on an Illumina MiSeq (Illumina, Inc., San Diego, CA, United States) in 2×300 bp run mode. Resulting sequence data was demultiplexed by the JGI and processed through JGI’s quality-control system which filters out known contaminant reads using the kmer filter in bbduk and also removes adaptor sequences (https://jgi.doe.gov/wp-content/uploads/2013/05/iTagger-methods.pdf). The quality-controlled sequence read files were downloaded and used for downstream analysis.

The JGI iTag SOP does not include the sequencing of negative controls or blanks. The JGI quality-controlled sequence reads generated for the ITS2 region were deposited at GenBank under BioProject ID <u>PRJNA667465</u> and for the 18S rRNA gene at <u>PRJNA667462</u>. Sequence reads are also available from the JGI Genome Portal (https://genome.jgi.doe.gov/portal/Popandseaspecies/Popandseaspecies.info.html).

### Sequence processing

Primers were removed using cutadapt (v. 2.1) (Martin, 2011). The resulting fastq files were analyzed in R (v. 4.0.2) using DADA2 (v. 1.12.1), phyloseq (v. 1.32.0), vegan (v. 2.5-6), microbiome (v. 1.10.0), ecodist (v. 2.0.5), EcoUtils (v. 0.1), DESeq2 (v. 1.28.1), ggplot2 (v. 3.3.2), tidyverse (v. 1.3.0) and many other R packages (Lahti & Shetty; Hothorn *et al.*, 2006, 2008; Dray & Dufour, 2007; Goslee & Urban, 2007; Wickham, 2007, 2016; Sarkar, 2008; Morgan *et al.*, 2009; Zeileis & Croissant, 2010; Eddelbuettel, 2013; Lawrence *et al.*, 2013; McMurdie & Holmes, 2013; Love *et al.*, 2014; Neuwirth, 2014; Xie, 2014; Huber *et al.*, 2015; Ritchie *et al.*, 2015; Callahan *et al.*, 2016; Elzhov *et al.*, 2016; Baselga *et al.*, 2018; Becker, 2018; Chen, 2018; Garnier, 2018; Hijmans, 2019; Oksanen *et al.*, 2019; Simpson, 2019; Wickham *et al.*, 2019; Allaire *et al.*, 2020; Bass *et al.*, 2020; Harrell *et al.*, 2020; Ogle *et al.*, 2020; Pedersen, 2020; Robinson & Hayes, 2020; Salazar, 2020; Sprockett, 2020; Therneau, 2020; Wickham & Seidel, 2020; Yu, 2020). For a detailed walkthrough of the following analysis using R, see the R-markdown summary file (Ettinger, 2020).

Prior to denoising in DADA2, reads were truncated at the first quality score of 2 and reads with an expected error greater than 2 were removed. Reads were then denoised and merged to generate tables of amplicon sequence variants (ASVs) using DADA2. Prior to downstream analyses, chimeric sequences were identified and removed from tables using removeBimeraDenovo (12.62% of sequences for ITS2 region, 4.53% of sequences for 18S rRNA gene). Taxonomy was inferred using the RDP Naive Bayesian Classifier algorithm with a modified UNITE (v. 8.2 “all eukaryotes”) database for ITS2 region sequences and the SILVA (v. 138) database for 18S rRNA gene sequences resulting in 89,754 and 53,084 ASVs respectively (Wang *et al.*, 2007; Quast *et al.*, 2013; Yilmaz *et al.*, 2014; Abarenkov *et al.*, 2020). The UNITE database was modified to include a representative ITS2 region amplicon sequence for the host plant, *Z. marina* (KM051458.1) as was done previously in Ettinger and Eisen (2019). ASVs were then each given a unique name by giving each a number preceded by “ITS” or “18S” and then “SV” which stands for sequence variant (e.g., ITS_SV1, ITS_SV2, etc. and 18S_SV1, 18S_SV2, etc.).

Based on the results of Pauvert et al. (2019), ITS-x was not run on the ITS2 region ASVs. However, we removed all ASVs taxonomically assigned as non-fungal at the domain level (e.g., ASVs assigned to the host plant, *Z. marina*, other eukaryotic groups or with no domain level classification) from the ITS2 region ASV table prior to downstream analysis resulting in a final table of 5,089 ASVs representing 488 samples (n_leaf_ = 179, n_root_ = 173, n_sediment_ = 136). A total of 88 samples were dropped from the analysis either because they had no sequences after being processed through DADA2 or no remaining sequences after removing non-fungal ASVs.

For the 18S rRNA gene ASV table, we generated two different filtered datasets (1) a fungal only dataset and (2) a general eukaryotic dataset. For (1), we removed all non-fungal ASVs from the 18S rRNA gene ASV table prior to downstream analysis of the fungi in this dataset resulting in a table of 1,216 fungal ASVs representing 409 samples (n_leaf_ = 146, n_root_ = 144, n_sediment_ = 119). A total of 167 samples were dropped from the analysis either because they had no sequences after being processed through DADA2 or because they had no remaining sequences after removing all ASVs classified as non-fungal. For (2), we removed ASVs taxonomically classified as non-eukaryotic and also as being from embryophytes (e.g. *Z. marina*) from the 18S rRNA gene ASV table resulting in a table of 36,582 eukaryotic ASVs representing 556 samples (n_leaf_ = 187, n_root_ = 187, n_sediment_ = 182). A total of 20 samples were dropped from the analysis either because they had no sequences after being processed through DADA2 or no remaining sequences after filtering ASVs.

### Sequence analysis and visualization

We utilized raw read counts, proportions, centered log-ratio, or Hellinger transformations on the data as appropriate when performing statistics and generating visualizations. Centered log-ratio and Hellinger transformations were performed using the transform function in the microbiome R package. Centered log-ratio (clr) values are scale-invariant such that the same ratio is obtained regardless of differences in read counts and thus were suggested as appropriate transformations for microbiome analysis by Gloor et al. (2017). When calculating abundance-occupancy curves, we used rarefy_even_depth in the phyloseq R package to subset to 1000 and 100 reads without replacement respectively for the ITS2 region and 18S ASV tables following the code in Shade and Stopnisek (2019).

To assess alpha (i.e. within sample) diversity between sample types (leaf, root, and sediment), the Shannon index of samples were calculated on ASV tables containing raw read counts using the estimate_richness function in the phyloseq R package. Raw read counts were used instead of normalizing the data by rarefying, as this kind of subsampling has been shown to be statistically inappropriate (McMurdie & Holmes, 2014). To assess alpha diversity across each of the 16 collection sites (Table S1) and across oceans, we first split the dataset into different sample types (leaf, root, and sediment) and then for each sample type, we calculated the Shannon index of samples. Kruskal–Wallis tests with 9,999 permutations were used to test for significant differences in alpha diversity across comparisons (sample type, site or ocean). For comparisons in which the Kruskal–Wallis test resulted in a rejected null hypothesis (*p* < 0.05), Bonferroni corrected *post hoc* Dunn tests were performed.

To assess beta (i.e. between-sample) diversity, we calculated several ecological metrics (Bray-Curtis, Aitchinson, Hellinger) using the ordinate function in phyloseq and visualized them using principal coordinates analysis. The Bray-Curtis dissimilarity is a widely used ecological metric in microbial analyses which calculates the compositional dissimilarity between samples (Bray *et al.*, 1957). The Aitchison distance, which is the Euclidean distance of clr transformed samples, is thought to be better than Bray-Curtis dissimilarity, because it is more stable to subsetting the data, and is also a true linear distance (Aitchison *et al.*, 2000; Gloor *et al.*, 2017). The Hellinger distance, which is the Euclidean distance of Hellinger transformed data, is based on differences in the proportions of taxa and is thought to be a more ecologically relevant representation of the composition of taxa between samples in comparison to Bray-Curtis dissimilarity which is biased towards abundant taxa (Rao, 1997; Legendre & Gallagher, 2001).

To test for significant differences in mean centroids between categories of interest (i.e. sample type, site, ocean) for each ecological metric (Bray-Curtis, Aitchinson, Hellinger), we performed permutational manovas (PERMANOVAs) with 9,999 permutations and to account for multiple comparisons, we adjusted *p*-values using the Bonferroni correction (Anderson, 2001). We also tested for significant differences in mean dispersions between different categories of interest using the betadisper and permutest functions from the vegan package in R with 9,999 permutations. The *post hoc* Tukey’s honest significant difference (HSD) test was performed on betadisper results that resulted in a rejected null hypothesis (*p* < 0.05), to identify which categories had mean dispersions that were significantly different.

To test for correlations between the community distances (Bray-Curtis, Hellinger) and geographic distances between samples, we first subset the data by ocean and sample type and then calculated the geographical distances between samples using the Haversine formula which accounts for the spherical nature of Earth using the distm function in the geosphere R package. Then we performed Mantel tests using 9,999 permutations and generated Mantel correlograms using the mantel and mantel.correlog functions in the vegan R package. To further support Mantel test results, we performed multiple regression on distance matrices (MRM) between community distances and geographic distances using 9,999 permutations via the MRM function in the ecodist R package. The code to perform distance-decay analyses was adapted from Wainwright et al (2019b).

To visualize global fungal community composition across sample types (leaf, root, and sediment), we transformed raw read counts to proportions and collapsed ASVs into taxonomic orders using the tax_glom function in phyloseq and then removed orders with a mean proportion of less than one percent. This threshold was chosen to better visualize only the most abundant orders, while also removing rare orders to avoid possible false positives during statistical analysis. The average relative abundance of taxonomic orders was compared between sample types using Bonferroni corrected Kruskal–Wallis tests in R and Bonferroni corrected *post hoc* Dunn tests were performed for orders where the Kruskal–Wallis test resulted in a rejected null hypothesis, to identify which sample type comparisons for each taxonomic order were significantly different.

To examine the contribution of specific ASVs to fungal community composition, we used the DESeq2 R package on the raw read counts to examine the log_2_fold change (differential abundance) of ASVs across sample types (leaf, root, sediment) in both datasets. We then visualized the global distribution of ASVs found to have significantly different differential abundances (Bonferroni corrected *p* < 0.05). To do this, we transformed the raw read counts to proportions and then subset each dataset to only include the single ASV of interest using prune_taxa in the phyloseq R package.

A core microbial community is usually defined as taxa that occur above an arbitrary detection threshold (e.g. greater than 1% relative abundance) and also above an arbitrary occupancy threshold (e.g. from 30% in Ainsworth et al. (2015) to 95% in Huse et al. (2012)). In an attempt to define “common” core leaf, root and sediment mycobiomes (“common” as defined in Risely (2020)), we used a more standardized approach by building abundance-occupancy curves and then calculating the rank contribution of specific ASVs to beta diversity (Bray-Curtis) to identify putative core ASVs using code from Shade and Stopnisek (2019). ASVs were predicted to be in the core using the final percent increase in beta-diversity method described in Shade and Stopnisek (2019) with a final percent increase of equal or greater than 10%. We then fit the Sloan neutral model (Sloan *et al.*, 2007) to the abundance-occupancy curves using the code provided in Burns et al. (2016) to predict whether core taxa were selected for by the environment (e.g. by the host plant, *Z. marina*), dispersal-limited or neutrally selected.

To investigate the general composition of the eukaryotic community and assess what proportion of the eukaryotic community is taxonomically classified as fungal, we first transformed raw read counts from the 18S rRNA gene ASV table filtered to include all eukaryotes to proportions and collapsed ASVs into taxonomic phyla using the tax_glom function in phyloseq. For visualization purposes, we then removed phyla with a mean proportion of less than 0.1 percent. The average relative abundance of eukaryotic phyla was then calculated for each sample type (leaf, root, sediment).

To investigate possible functional roles of seagrass-associated fungi, FUNGuild (v. 1.1) was run on the taxonomic assignments of ASVs from both the ITS2 region and 18S rRNA gene datasets (Nguyen *et al.*, 2016). FUNGuild searches the taxonomic assignments at the genus level against an online Guilds database containing taxonomic keywords and functional metadata (e.g. trophic level, guild, etc.) and FUNGuild assignments are given confidence rankings of “Highly Probable”, “Probable” or “Possible”. To assess ecological guilds of high confidence, we first visualized all annotations that were ranked as “highly probable” in either dataset. We then investigated functional guilds that were assigned to only highly abundant ASVs in our data. To assess this, we subset both the ITS2 region and 18S rRNA gene ASV tables to include only ASVs with a mean abundance of greater than 0.1 percent and then visualized the data in R.

## Results

### Fungal alpha diversity differs between sites, tissues and oceans

The Shannon index was significantly different between sample types (K-W test, *p* < 0.001, Figure 1) for both the ITS2 region amplicon and 18S rRNA gene amplicon datasets. *Post hoc* Dunn tests of both datasets suggest that alpha diversity for leaves was consistently lower than that of the roots (*p* < 0.05). However, there were conflicting results for the sediment, with diversity being lower in the sediment than leaves and roots in the ITS2 region amplicons (*p* < 0.05) and diversity being higher in the sediment than leaves and roots in the 18S rRNA gene amplicons (*p* < 0.05). Alpha diversity for both datasets also was significantly different within each sample type across sites (K-W test, *p* < 0.001, Figure S1). This was driven by diversity being significantly different across some, but not all sites (Dunn, *p* < 0.05). Alpha diversity for leaves was significantly different across oceans for the ITS2 region amplicon dataset (K-W test, *p* = 0.0142), but was not significantly different for roots or sediment between oceans or for leaves, roots or sediment between oceans for the 18S rRNA gene amplicon dataset (*p* > 0.05).

**Figure 1.**
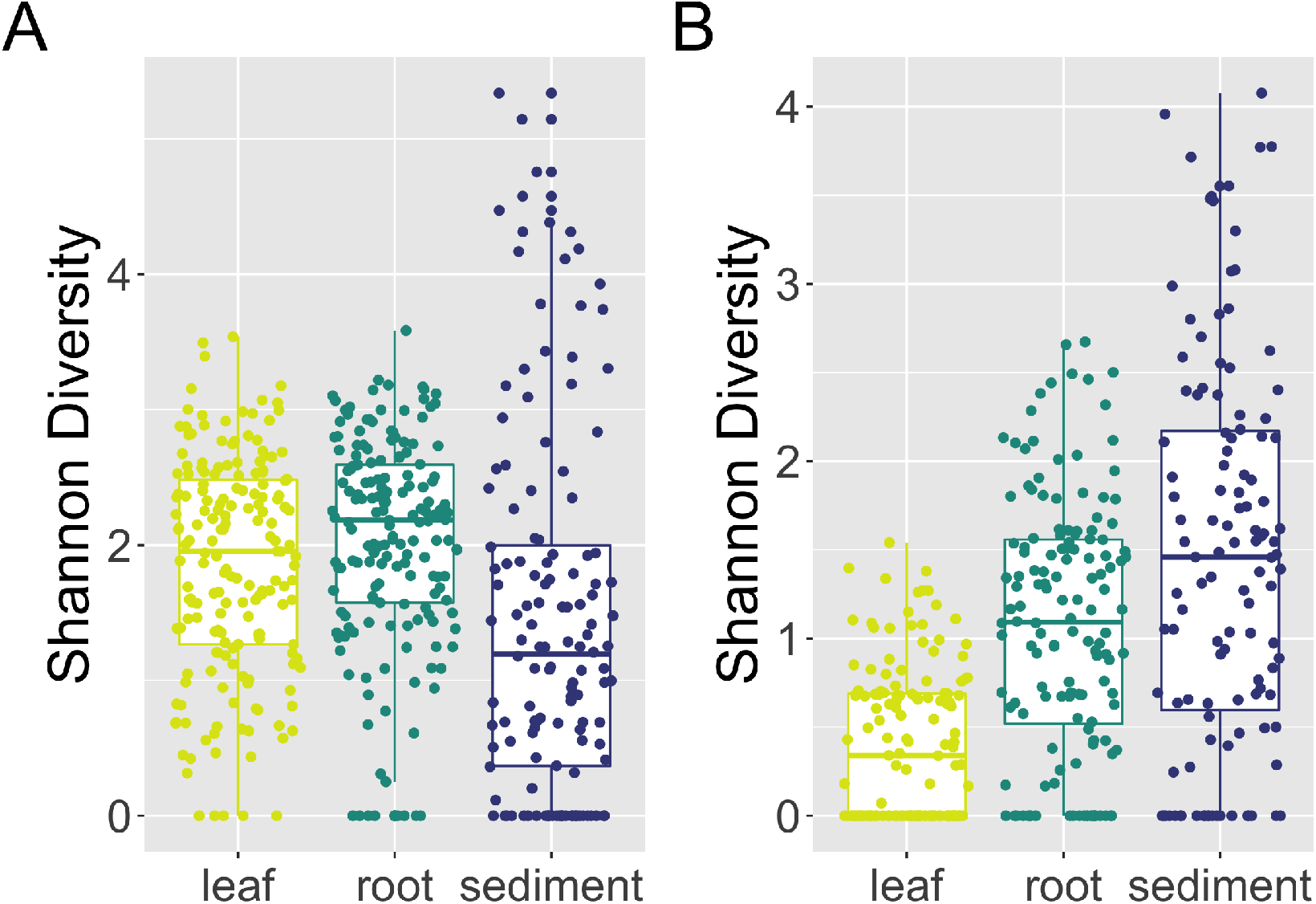
Within sample diversity varies across tissues. Boxplot visualizations of Shannon diversities for each sample type (leaf, root, sediment) based on (A) ITS2 region amplicon data and (B) 18S rRNA gene amplicon data.

### Fungal community structure differs across sites, tissues and oceans

Similar to alpha diversity, fungal beta diversity was significantly different for both datasets using all three ecological metrics (Bray-Curtis, Aitchinson, Hellinger) across sample types (PERMANOVA, *p* < 0.001, Figure 2), across sites (*p* < 0.001, Figure S2) and across oceans (*p* < 0.001, Figure S2). *Post hoc* pair-wise PERMANOVA tests using the ITS2 region amplicon data indicated significant differences in beta diversity across sample types (*p* < 0.001) and sites (*p* < 0.01). These results were generally consistent with the 18S rRNA gene sequence data which supported differences in community structure across sample types (*p* < 0.001) and across most, but not all, collection sites (*p* < 0.05).

**Figure 2.**
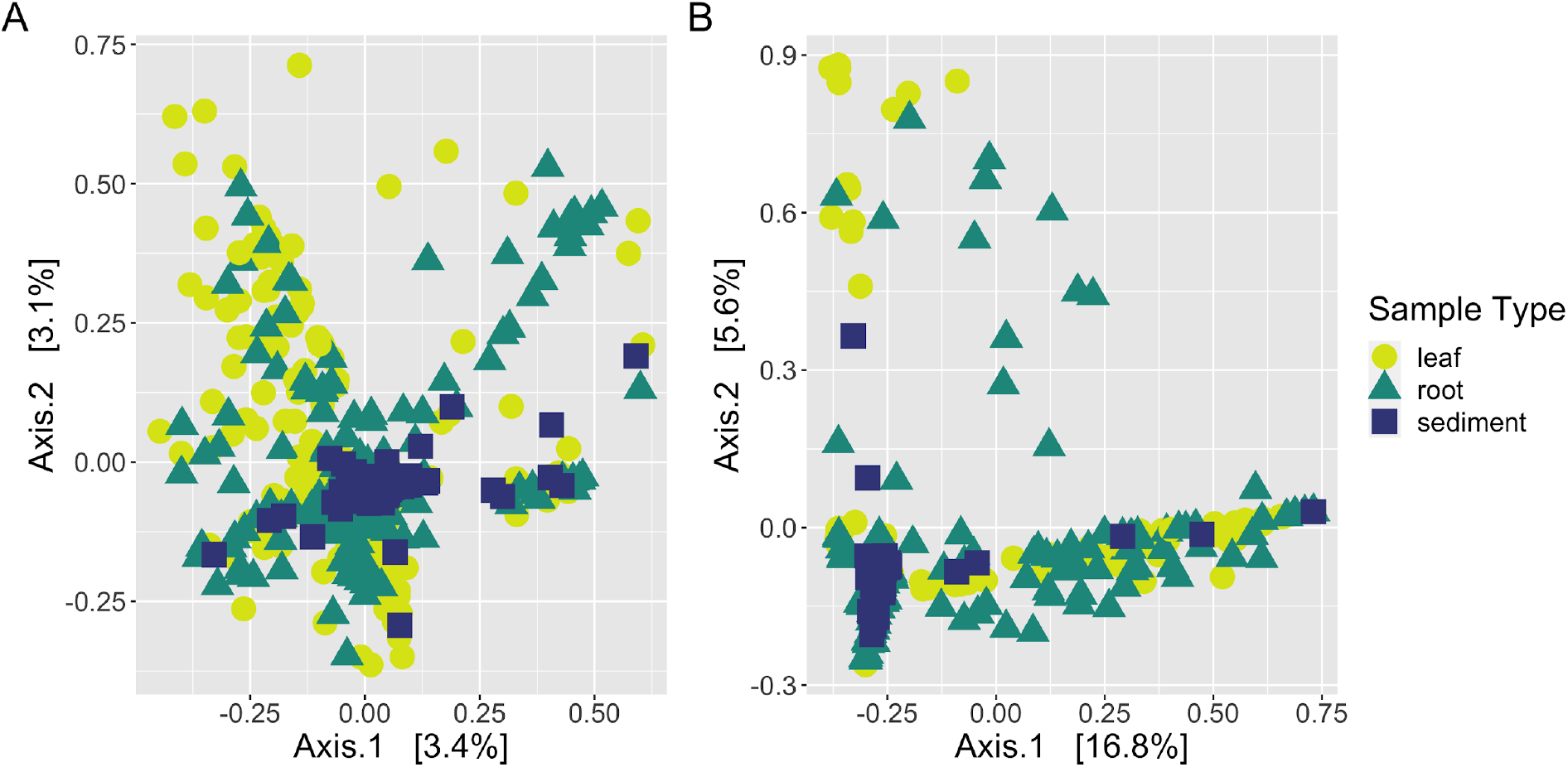
Community structure varies between tissues. Principal coordinates analysis (PCoA) visualization of Hellinger distances of fungal communities associated with leaves, roots, and sediment based on (A) ITS2 region amplicon data and (B) 18S rRNA gene amplicon data. Points in the ordination are colored and represented by shapes based on sample type: leaf (yellow circles), root (green triangles) or sediment (blue squares).

Within group variance (i.e. dispersion) also differed significantly for the ITS2 region amplicon data using all three beta diversity metrics across sample types (betadisper, *p* < 0.01) and sites (*p* < 0.01), but did not vary across oceans (*p* > 0.05). Mean dispersion between sites in the 18S rRNA gene data was not significant for two of the ecological metrics (Bray-Curtis: *p* = 0.79, Hellinger: *p* = 1) and similarly the mean dispersion between oceans was not significant for two of the ecological metrics (Bray-Curtis: *p* = 0.07, Aitchinson: *p* = 0.26). Mean dispersion was otherwise consistent with the significant results observed in the ITS2 region amplicon data. PERMANOVA results have been shown to confuse dispersion differences and centroid differences when not using a balanced design. Therefore, our results may indicate that either mean centroids, mean dispersions, or both are differing between sample types and sites here.

### Mantel tests suggest weak distance-decay relationships within oceans

Mantel tests indicated a small, but significant positive relationship between both metrics of community structure (Bray-Curtis, Hellinger) and geographic distance for leaves across the Pacfic Ocean for the ITS2 region and 18S rRNA gene amplicon datasets (*p* < 0.001, Figure 3A, Figure S3A, Table S2). This relationship was also detected for leaves across the Atlantic Ocean (*p* < 0.001, Figure 3B, Figure S3B, Table S2). Mantel correlograms suggest that this pattern is driven by sites with the closest proximity, such that sites closer together have more similar fungal communities compared to sites further away (Figure S4). In the Pacific Ocean, roots had consistently the strongest positive relationship with geographic distance for both the ITS2 region and 18S rRNA gene amplicon datasets (*p* < 0.001, Figure S3C, Figure S5A, TableS2). Interestingly, a much weaker, but still significant, positive relationship was observed for roots in the Atlantic Ocean (*p* < 0.001, Figure S3D, Figure S5B, Table S2). Sediment in the Pacific Ocean had the weakest relationship with distance with conflicting significance for the ITS2 region (Bray-Curtis: *p* < 0.001; Hellinger: *p* = 0.827) and only small, but still significant correlations for 18S rRNA gene amplicon datasets (Bray-Curtis: *p* = 0.031; Hellinger: *p* = 0.011, Figure S3E, Figure S5C, Table S2). In contrast in the Atlantic Ocean, sediment had much more robust positive relationships with geographic distance for both datasets (*p* < 0.001, Figure S3F, Figure S5D, Table S2). Multiple regression analyses further confirmed all significant patterns of observed distance-decay (*p* < 0.001).

**Figure 3.**
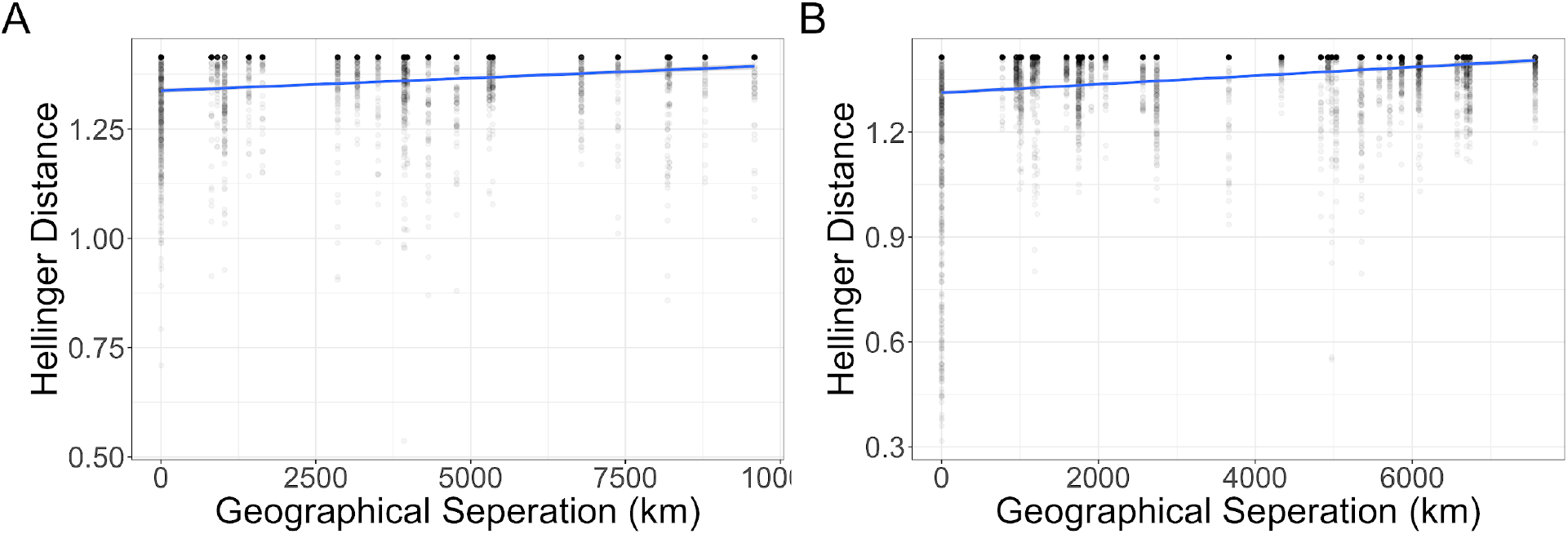
Mantel tests suggest a distance-decay relationship. Scatterplots depicting the weak, but significant positive distance–decay relationship between leaf fungal community beta diversity (Hellinger distance) using the ITS2 region amplicon data and geographical distance (km) between sites from the (A) Pacific Ocean, and (B) Atlantic Ocean.

### Mean taxonomic composition of the global mycobiome

The majority of taxonomic orders had mean relative abundances that were significantly different between sample types in both the ITS2 region amplicon and 18S rRNA gene amplicon datasets (K-W test, *p* < 0.01, Figure S6) and many of these were enriched on *Z. marina* tissues over rhizosphere sediment. In the ITS2 region amplicon data, Dothideales, Lobulomycetales, and unclassified Sordariomycetes were all enriched on both leaf and root tissues relative to sediment (Dunn, p < 0.01). Polyporales, Helotiales, Hypocreales, Capnodiales and Malasseziales were all in higher abundance on leaves (p < 0.001). Unclassified Ascomycota had increased relative abundance on roots (p < 0.001). Fungi that were unable to be classified to the phylum level were more abundant on roots and in rhizosphere sediment relative to leaves (p < 0.001). Comparatively, in the 18S rRNA gene data, Saccharomycetales were enriched on the leaves (p < 0.001) and unclassified Chytridiomycota and unclassified Sordariomycetes were in greater abundance on roots (p < 0.001). We attribute differences between the two datasets to the use of different primer sets which each have their own biases, as well as the different reference databases used for each amplicon to assign taxonomy.

### Global core leaf, root and sediment mycobiomes

We utilized abundance-occupancy distributions of ASVs to infer global *Z. marina* leaf, root and rhizosphere sediment core mycobiomes based on ASV rank contributions to beta diversity. A total of 14, 15, and 60 ASVs were predicted as being in the leaf, root, and sediment cores respectively based on the ITS2 region amplicon data (Figure 4A, Table S3). Four ASVs overlapped across all three cores; this included generalist fungi with widespread distributions like *Cladosporium sp.* and *Malassezia restricta* (Amend, 2014; Ettinger & Eisen, 2020). Interestingly, only one ASV was shared between leaf and root cores, *Saccharomyces paradoxus* (ITS_SV260). The leaf core was dominated by unclassified Capnodiales spp., while the root core was dominated by unclassified Sordariomycetes spp. The sediment core was more diverse, but was composed mostly of Ascomycota, particularly members in the Pleosporales and Agaricales.

**Figure 4.**
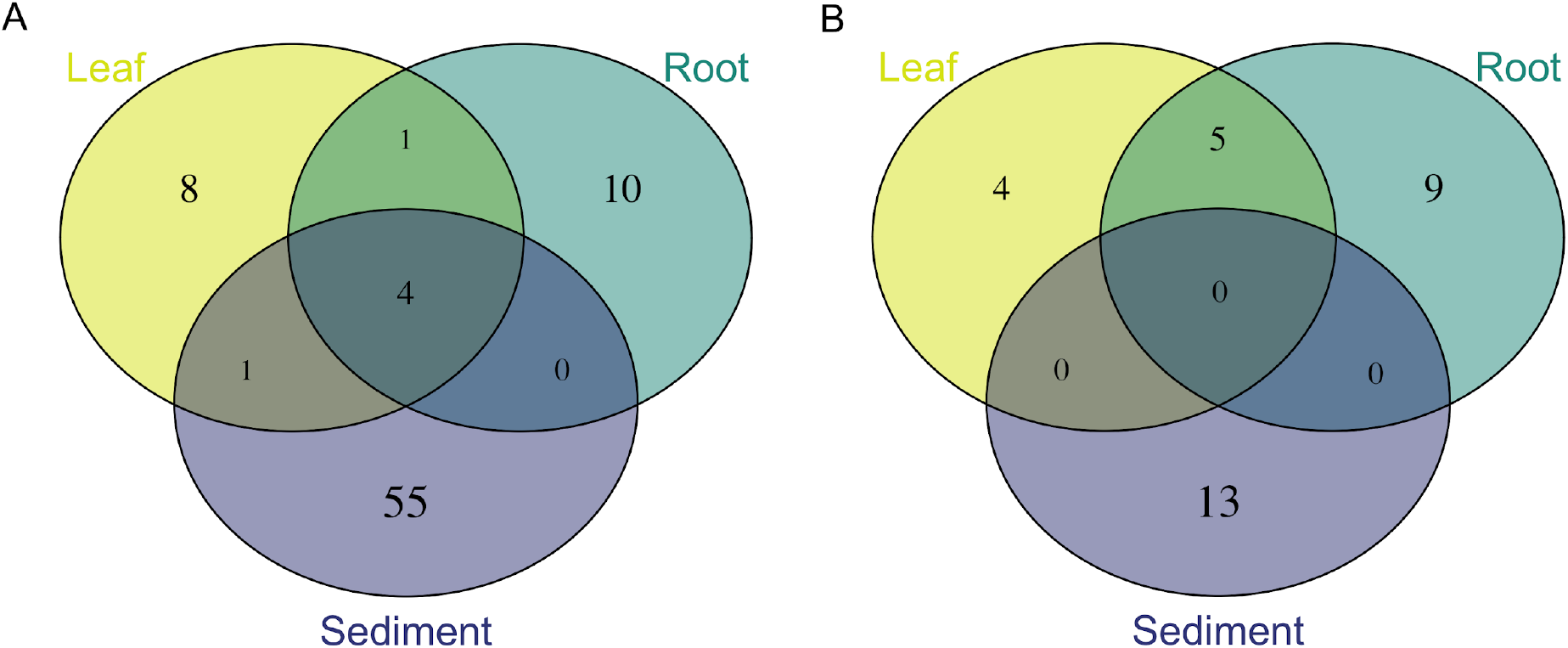
Overlap between predicted core mycobiomes of individual *Z. marina* tissues. Venn diagrams representing shared core ASVs as defined by abundance-occupancy distributions for each sample type (leaf, root, sediment) for (A) ITS2 region amplicon data, and (B) 18S rRNA gene amplicon data.

Smaller core mycobiomes were predicted from the 18S rRNA gene amplicon data with only 9, 14, and 13 ASVs placed in the leaf, root, and sediment cores, and no ASVs overlapped between the three cores (Figure 4B, Table S4). However, five ASVs were shared between leaf and root cores, three belonging to unclassified Chytridiomycota lineages, a *Saccharomyces* sp. and an unclassified Sordariomycetes sp. A total of four ASVs were unique to the leaf core which was dominated by unclassified Chytridiomycota lineages and nine ASVs were unique to the root core which was predominantly comprised of unclassified Sordariomycetes spp. and unclassified Chytridiomycota lineages (including Lobulomycetaceae spp.). The sediment core was mostly made of Saccharomycetales lineages and Chytridiomycota lineages.

### Neutral models to predict ASV selection

We applied Sloan neutral models to investigate if core ASVs are selected for by *Z. marina*, assembled through stochastic or deterministic processes (Sloan *et al.*, 2007; Burns *et al.*, 2016). ASVs that fall above the neutral model prediction appear in higher occupancy than would be predicted based on their relative abundance and are thus, thought to be selected for by the plant environment. ASVs that fall below the neutral model prediction have higher relative abundance than would be predicted based on their occupancy and are thus, thought to be either selected-against by the plant host or dispersal-limited. For the ITS2 region abundance-occupancy distributions, 2.9%, 4.84%, and 3.74% of all ASVs fell above/below the neutral model prediction for leaves, roots and sediment respectively (Figure 5). While for the 18S rRNA gene abundance-occupancy distributions, 7.5%, 6.44%, and 2.16% of all ASVs deviated from the neutral model (Figure S7). Further, looking at deviations from the neutral model for ASVs predicted to be in the core mycobiome allows insight into the role of *Z. marina* in core assembly. We found that of the core leaf, core root and core sediment ASVs several were predicted to be plant-selected (n_leaf_ = 6, n_root_= 7, n_sediment_ = 40), only a few were selected-against or dispersal-limited (n_leaf_ = 1, n_root_= 3, n_sediment_ = 4), and most were neutrally selected (n_leaf_ = 16, n_root_= 19, n_sediment_ = 29).

**Figure 5.**
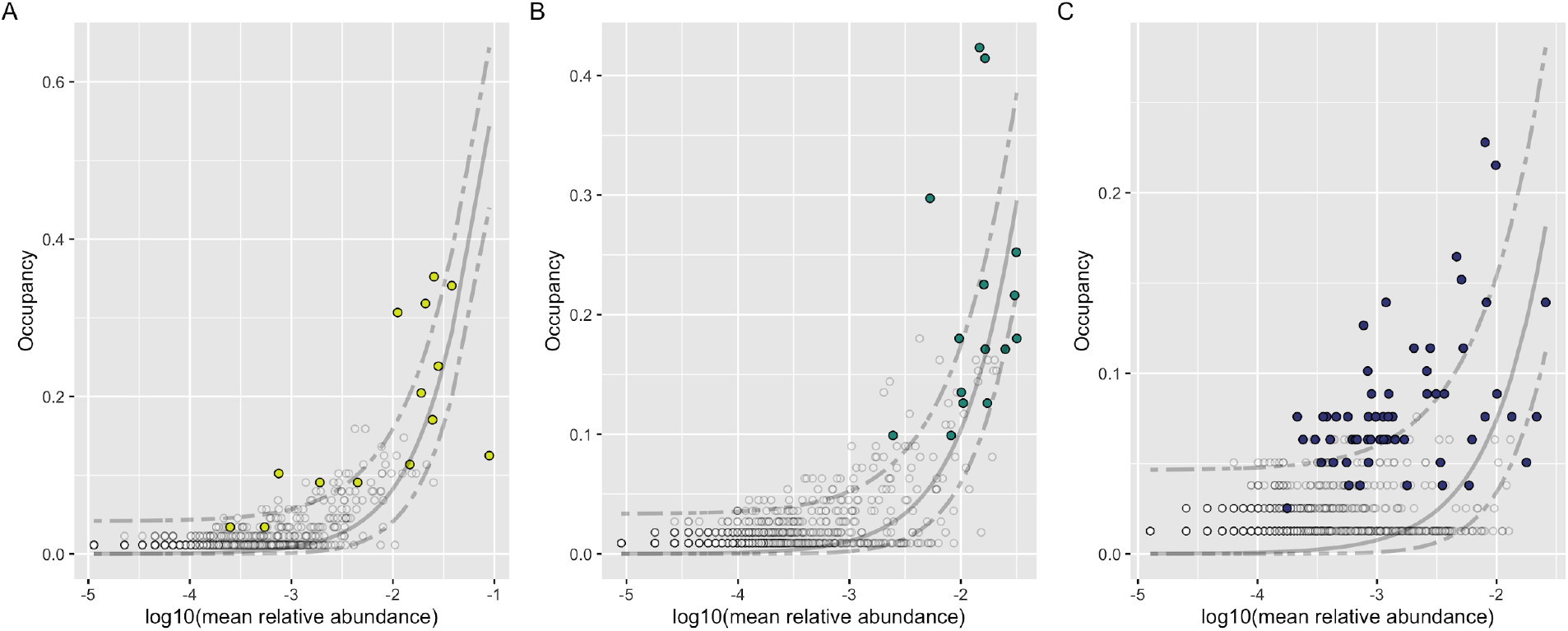
Abundance-occupancy distributions reveal core mycobiomes. Abundance-occupancy distributions were used to define core members of the (A) leaf, (B) root and (C) sediment mycobiomes for the ITS2 region amplicon data. Each point represents an ASV with predicted core members indicated by a color (leaf = yellow, root = green, sediment = blue) and non-core ASVs in white. Ranked ASVs were predicted to be in the core based on a final percent increase of equal or greater than 10%. The solid line represents the fit of the neutral model, and the dashed line is 95% confidence around the model prediction. ASVs above the neutral model are predicted to be selected for by the environment (e.g. by the host plant, *Z. marina*), and those below the model are predicted to be selected-against or dispersal-limited.

Generally the neutral models had poor fits for both the ITS2 region (leaf: *R*^*2*^ = 0.31; root: *R*^*2*^ = 0.44; sediment: *R*^*2*^ = −0.76), and 18S rRNA gene datasets (leaf: *R*^*2*^ = 0.49; root: *R*^*2*^ = 0.50; sediment: *R*^*2*^ = 0.08), with the sediment curves having the worst fit to the neutral model. This could potentially be attributed to the low predicted migration rates for both the ITS2 region (leaf: *m* = 0.001; root: *m* = 0.002; sediment: *m* = 0.001) and 18S rRNA gene datasets (leaf: *m* = 0.002; root: *m* = 0.014; sediment: *m* = 0.014). These values are consistent with other studies of fungi that used neutral models on abundance-occupancy curves of fungi (Stopnisek & Shade) and may be reflective of dispersal limitation playing a stronger role in fungal assembly than bacterial community assembly (Talbot *et al.*, 2014; Tedersoo *et al.*, 2014; Gumiere *et al.*, 2016).

### Global distribution of differentially abundant ASVs

To investigate variation in fungal community composition at greater taxonomic resolution, we used DESeq2 to identify ASVs whose abundance differed across sample types (Figure S8 and S9). The greatest number of differentially abundant ASVs was observed between the roots and sediment, with fourteen ITS2 region ASVs and four 18S rRNA gene ASVs (Wald test, *p* < 0.01). This was closely followed by differentially abundant ASVs between leaves and sediment, with twelve ITS2 region ASVs and two 18S rRNA gene ASVs (*p* < 0.05). The smallest number of differentially abundant ASVs was found between leaves and roots, with three ITS2 region ASVs (*p* < 0.01). We compared the differentially abundant ASVs to those predicted to be in the leaf, root and sediment core mycobiomes. We found fourteen ASVs that were both differentially abundant between sample types and present in at least one core mycobiome; of those fourteen, seven were also found to deviate from the neutral model (Table 1). We then examined the global distribution of the fourteen ASVs that were both differentially abundant across sample types and predicted to be in the *Z. marina* core mycobiome. For example, ITS_SV260 (*Saccharomyces paradoxus*) appears to be globally distributed, neutrally selected, and more abundant on leaves and roots than sediment (*p* < 0.001, Figure 6). In contrast, ITS_SV362 (*Lobulomyces* sp.) appears to be only found at one site, dispersal-limited, and is more abundant on leaves than sediment (*p* < 0.001, Figure 7).

**Table 1.** Predicted differentially abundant core ASVs. ASVs were ranked by abundance-occupancy distributions and then predicted to be in a core based on a final percent increase in beta-diversity of equal or greater than 10%. The Sloan neutral model was then applied to the abundance-occupancy distributions to identify ASVs that deviate such that ASVs above the neutral model are predicted to be selected for by the environment (e.g. by the host plant, *Z. marina*), and those below the model are predicted to be selected-against or dispersal-limited. Finally, DESeq2 was used to identify ASVs that were differentially abundant between pair-wise sample types (leaf, root, sediment). Here for each predicted core ASV that was also differentially abundant for at least one pairwise comparison, we report the ASV, the core it was predicted to be a member of (leaf, root or sediment), whether the ASV deviated from the neutral model (above, below or none), the significant pairwise differential abundance comparisons (e.g. root > sediment means that the ASV was in significantly higher abundance when associated with roots than with sediment), and the taxonomy of the ASV.

| ASV | Core prediction | Neutral model<br>deviations | Significant DESeq2<br>comparisons | Taxonomy |
| --- | --- | --- | --- | --- |
| ITS_SV52 | leaf, root,<br>sediment | above, above, none | root > sediment | *Mycosphaerella tassiana* |
| ITS_SV60 | root | below | leaf > sediment; root ><br>sediment | Unclassified Sordariomycetes sp. |
| ITS_SV125 | root | none | root > sediment | Unclassified Ascomycota sp. |
| ITS_SV234 | root | none | leaf > sediment; root ><br>sediment | Unclassified Sordariomycetes sp. |
| **ITS_SV260** | leaf, root | none, none | leaf > sediment; root > sediment | *Saccharomyces paradoxus* |
| **ITS_SV362** | leaf | below | leaf > sediment; root > sediment | *Lobulomyces* sp. |
| **ITS_SV426** | leaf, sediment | none, none | leaf > sediment; root > sediment | *Saccharomyces* sp. |
| **ITS_SV497** | root | below | leaf > sediment; root > sediment | Unclassified Sordariomycetes sp. |
| **ITS_SV679** | sediment | above | root > leaf; root > leaf | *Pseudeurotium bakeri* |
| **ITS_SV1045** | root | above | leaf > sediment; root > sediment | *Hortaea werneckii* |
| **18S_SV756** | leaf, root | none, none | root > sediment | Unclassified Chytridiomycetes sp. |
| **18S_SV928** | leaf, root | above, above | leaf > sediment; root > sediment | *Saccharomyces* sp. |
| **18S_SV968** | leaf | none | leaf > sediment; root > sediment | Unclassified Lobulomycetaceae sp. |
| **18S_SV1977** | root | none | root > sediment | *Chytridium* sp. |

**Figure 6.**
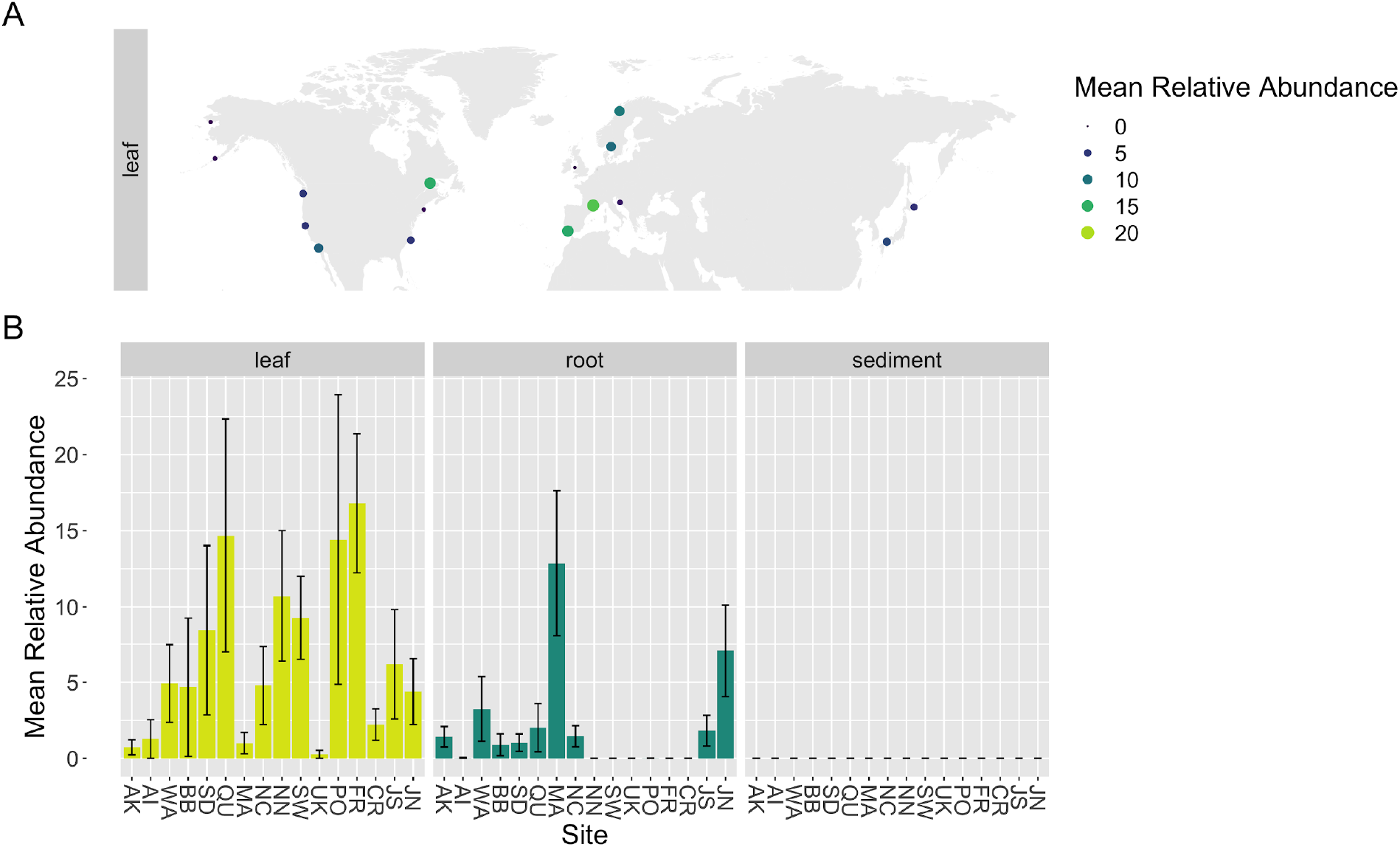
Example of differentially abundant neutrally selected core ASV. Here we show the global distribution of ITS_SV260, an ASV predicted to be a neutrally selected member of the core mycobiomes of both leaves and roots and also differentially abundant between leaves and sediment (*p* < 0.001), and roots and sediment (*p* < 0.001) using DESeq2. In (A) we plot the mean relative abundance of ITS_SV260 at each site on leaves on a global map, and in (B) we plot the mean relative abundance of ITS_SV260 on leaves, roots and sediment across sites, with the standard error of the mean represented by error bars and bars colored by sample type.

**Figure 7.**
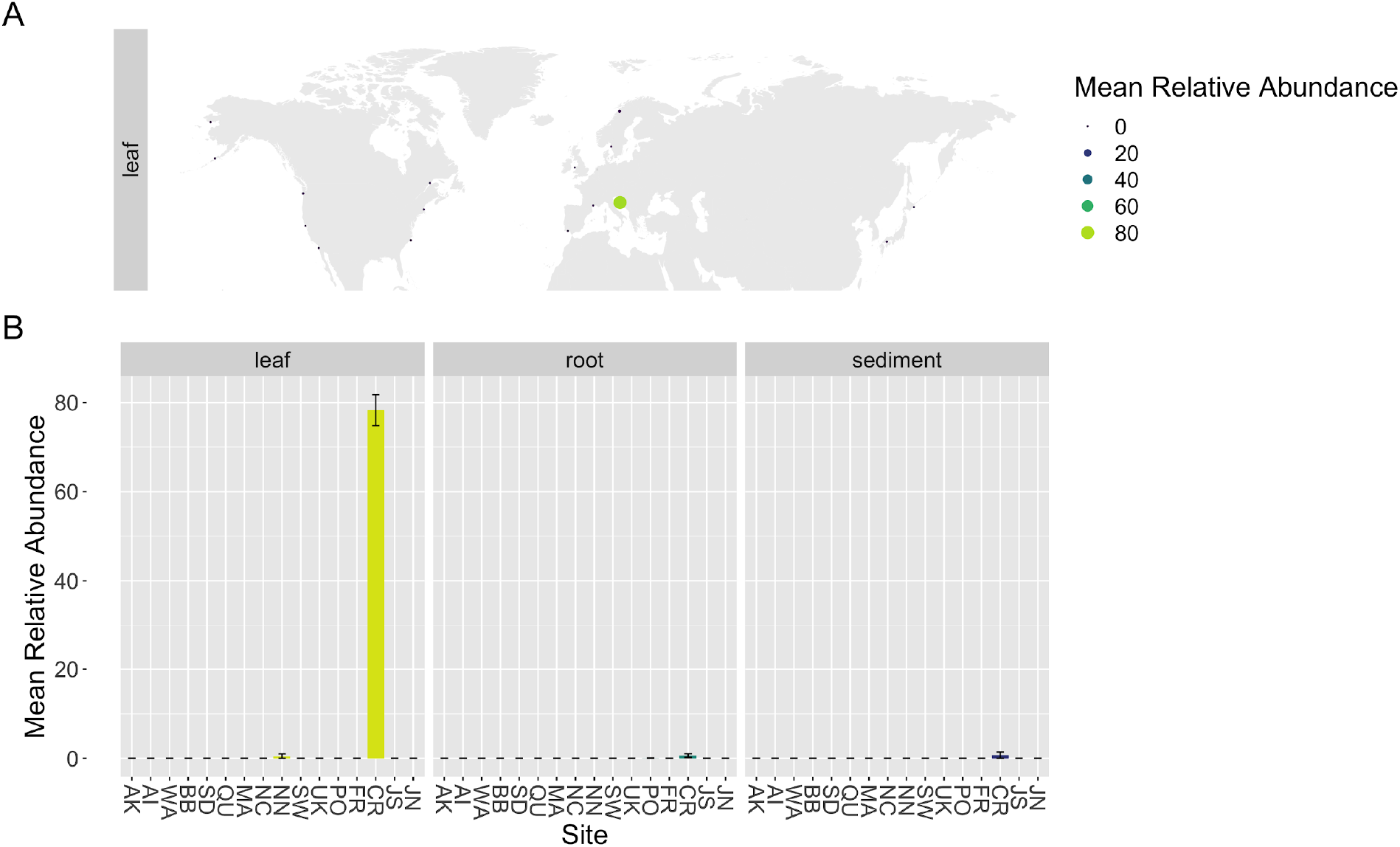
Example of differentially abundant dispersal-limited core ASV. Here we show the global distribution of ITS_SV362, an ASV predicted to be dispersal-limited and a member of the core mycobiomes of leaves and also differentially abundant between leaves and sediment (*p* < 0.001), and roots and sediment (*p* < 0.001) using DESeq2. In (A) we plot the mean relative abundance of ITS_SV362 at each site on leaves on a global map, and in (B) we plot the mean relative abundance of ITS_SV362 on leaves, roots and sediment across sites, with the standard error of the mean represented by error bars and bars colored by sample type.

### Fungi are only a small portion of *Z. marina* associated eukaryotic community

Fungal sequences made up only a tiny portion of the entire epiphytic eukaryotic community associated with *Z. marina* with a mean relative abundance on leaves of 0.50 ± 2.12%, roots of 0.12 ± 0.36%, and sediment of 0.23 ± 0.67% in the 18S rRNA gene dataset (Figure S10). The leaf eukaryotic community was generally dominated by diatoms, the root community by both diatoms and Peronosporomycetes (i.e. oomycetes) and the sediment community by both diatoms and dinoflagellates.

### Many ASVs have no predicted functional guild

Although FUNGuild was able to predict the functional guild and trophic mode of 78.62% of ASVs in the ITS2 region amplicon dataset, only 10.12% of ASVs had predictions at a confidence of “Highly Probable”. The most abundant functional guilds assigned at this confidence level included wood saprotroph, ectomycorrhizal, lichenized, endophyte, plant pathogen-wood saprotroph, and fungal parasite (Figure S11). Comparatively, FUNGuild was only able to predict functions for 35.31% of the ASVs in the 18S rRNA gene dataset and only 3.4% of those ASVs had “Highly Probable” predictions. Generally, the most abundant functional guilds at this confidence level were consistent with those in the ITS2 region dataset and included wood saprotroph, ectomycorrhizal, and plant pathogen (Figure S12). When we further investigated the predicted trophic modes of only the most abundant ASVs (mean relative abundance greater than 0.1 percent) in both the ITS2 region amplicon and 18S rRNA amplicon datasets, 38.38% and 63.5% of these ASVs respectively were unable to be assigned a function, a testament to how little we know about the functional roles of fungi in this system.

## Discussion

Our study of the *Zostera marina* mycobiome provides insight into the global distribution of host-associated fungi in the marine environment and highlights the need for future studies of marine fungal community dynamics and function. We found that the fungal community was different between sites globally and observed a small, but significant pattern of distance-decay for the *Z. marina* mycobiome. We defined a small core mycobiome for leaves, roots and sediment dominated by Sordariomycetes spp., Chytridiomycota lineages (including Lobulomycetaceae spp.), Capnodiales spp. and. Many differentially abundant core ASVs (e.g. *Lobulomyces* sp.) were only found at one or a few locations (e.g. possibly due to local adaptation, dispersal limitation or seasonal bloom events), while others (e.g. *Saccharomyces* sp.) were more ubiquitous across all locations suggesting a true global distribution and selection by the plant itself. Additionally, between the observed pattern of distance-decay, the shape of the fungal abundance-occupancy curves and the poor fit of the Sloan neutral model, it appears that, although affected by both stochastic and deterministic processes, the mycobiome of *Z. marina* may be more affected by deterministic factors (e.g. environmental filtering, host genetics, dispersal limitation) than perhaps expected. Finally, we found a large portion of ASVs were unable to be classified taxonomically and most ASVs were not able to be assigned a predicted functional guild, further highlighting how little we know about seagrass-associated fungi.

This study is the first to characterize the *Zostera marina* mycobiome across its full biogeographic distribution using culture-independent methods. We observed significant differences in alpha diversity both across seagrass tissues and across collection sites. From the 18S rRNA data, we observed that the alpha diversity of the sediment is more diverse than *Z. marina* tissues which is consistent with previous seagrass work (Wainwright *et al.*, 2019b; Hurtado-McCormick *et al.*, 2019; Ettinger & Eisen, 2019). However, across both datasets, we also found that leaves had a lower alpha diversity than roots, which is not consistent with our previous study of *Z. marina* (Ettinger & Eisen, 2019). This may be due to the different primer sets used in both studies as the use of different sequencing primers has been shown to have drastic effects on the results of mycobiome studies (Frau *et al.*, 2019). Primer bias and different reference databases may additionally explain some of the variation we found within this study between our two different amplicon datasets. This is not the first study to observe alpha diversity varying across sites. Previous seagrass work found that alpha diversity varied between sites (Wainwright *et al.*, 2019b; Hurtado-McCormick *et al.*, 2019), while other work found no differences in alpha diversity between sites (Trevathan-Tackett *et al.*, 2020).

In addition to differences in alpha diversity, we also observed differences in fungal community structure across tissues and sites. Differences in fungal beta diversity across sites and tissues has been reported previously for seagrasses (Bengtsson *et al.*, 2017; Wainwright *et al.*, 2018, 2019b; Hurtado-McCormick *et al.*, 2019; Ettinger & Eisen, 2019; Trevathan-Tackett *et al.*, 2020). Seasonal differences in fungal colonization of seagrasses have been observed previously and are likely contributing to the variation observed here between sites (Mata & Cebrián, 2013). The global planktonic marine fungal community has been found to cluster by ocean (Hassett *et al.*, 2020) and differences between oceans were observed here as well. However, site-to-site variation was a stronger factor driving differences suggesting that environmental or host plant filtering may play a critical role in assembling the fungal community associated with *Z. marina*.

It has long been thought that there are few barriers to fungal dispersal (Hyde *et al.*, 1998; Finlay, 2002; Fenchel & Finlay, 2004; Cox *et al.*, 2016). However, not every fungus is everywhere (Peay *et al.*, 2010), and there is increasing evidence for rampant environmental filtering and barriers to fungal dispersal for host-associated fungi in the marine ecosystem (Wainwright *et al.*, 2018, 2019a,b). The importance of biogeography for seagrass-associated fungal community structure can be seen in our observation of a small, but significant positive distance-decay relationship between geographic distance and community structure. This relationship suggests that sites closer together have more similar fungal communities compared to sites that are more distant from each other. Previously, similar distance-decay relationships were found present for other seagrass-associated fungal communities including with the seagrass, *Enhalus acoroides*, in Singapore and Peninsular Malaysia (2019b) and the seagrass, *Syringodium isoetifolium,* along Wallace’s line (Wainwright *et al.*, 2018).

The observed positive relationship between community structure and geographic distance is likely driven by a combination of factors including dispersal limitation, environmental filtering caused by local habitat differences and priority effects. Another factor that might be driving site-specific fungal community composition is host plant genetics. Host plant genotype has been found in other studies to strongly correlate with leaf fungal communities (Bálint *et al.*, 2013; Hunter *et al.*, 2015; Sapkota *et al.*, 2015). The natural dispersal distance of *Z. marina* is thought to be less than 150 km and there is some evidence of poor connectivity between locations and rampant inbreeding within locations (Olsen *et al.*, 2004; Muñiz-Salazar *et al.*, 2005; Campanella *et al.*, 2010; Ort *et al.*, 2012). Given the strong population structure and weak dispersal of *Z. marina,* variation in *Z. marina* genotypes could be playing a role in structuring the fungal community differences observed here. However, it should be noted that Wainwright et al. failed to find a correlation between *S. isoetifolium* genetics and fungal community composition in their study (2018). Regardless, there is growing evidence that seagrass-associated fungal communities are more similar at closer distances, and future work should look for correlations between *Z. marina* genetics, *Z. marina* dispersal and the fungal community.

Even though there were site-to-site differences in community structure, the global mycobiome of *Z. marina* was generally composed of members of taxonomic orders previously observed to associate with *Z. marina* and other seagrass species using culture-independent methods (Wainwright *et al.*, 2018, 2019b; Hurtado-McCormick *et al.*, 2019; Ettinger & Eisen, 2019, 2020; Trevathan-Tackett *et al.*, 2020). This diversity is also in line with cultivation efforts which have found Eurotiomycetes, Dothideomycetes, and Sordariomycetes to be the main classes of fungi associated with seagrasses (Sakayaroj *et al.*, 2010; Supaphon *et al.*, 2017; Ettinger & Eisen, 2020). Altogether our results are consistent with previous reports that the seagrass mycobiome is comprised of many ASVs, including many Chytridiomycota lineages, for which a specific taxonomic assignment cannot be made based on current datasets that are biased towards terrestrial fungi (Wainwright *et al.*, 2019b; Ettinger & Eisen, 2019; Trevathan-Tackett *et al.*, 2020). Likely contributing to this bias, only a few lineages of marine Chytridiomycota have been described using culture-based methods (Jones *et al.*, 2015, 2019) despite their dominance in DNA-based surveys of the marine environment (Hassett *et al.*, 2017, 2020; Ettinger & Eisen, 2019). The inability to taxonomically classify fungal sequences is a persistent problem for studies of the marine environment generally, and again serves to highlight the need for additional descriptive studies of these understudied marine organisms (Comeau *et al.*, 2016; Nagano *et al.*, 2017; Picard, 2017; Hassett *et al.*, 2017, 2020).

Despite being unable to taxonomically classify many *Z. marina* associated fungal sequences, we were still able to identify a small “common” core community associated with *Z. marina* tissues, with only a few ASVs unique to or shared between leaves and roots (Figure 4). Previously, Trevathan-Tackett et al. (2020) were able to identify a small core of eight fungal operational-taxonomic units (OTUs) associated with the leaves of *Zostera muelleri,* while Hurtado-McCormick (2019) were unable to identify a core fungal community on *Z. muelleri* leaves. The *Z. marina* core leaf and root mycobiomes were dominated by Sordariomycetes spp., Chytridiomycota lineages (including Lobulomycetaceae spp. which have previously been seen to dominate on this species (Ettinger & Eisen, 2019)) and Saccharomyces spp. (Table 1, Table S3, Table S4). Sordariomycetes were also found to dominate the core leaf mycobiome in Trevathan-Tackett et al. (2020). Only four ASVs overlapped between the core communities for leaves, root and sediment and these ASVs largely were assigned to known ubiquitous marine generalists (e.g. *Cladosporium* (Ettinger & Eisen, 2020), *Malassezia* (Amend, 2014)).

The expected shape of a microbial abundance-occupancy distribution is an ‘S’, with abundant taxa having the highest occupancies and rare taxa having the lowest occupancies (Shade & Stopnisek, 2019). However, the abundance-occupancy distributions for our data here do not have this shape. One possible reason for this deviation is an increased incidence of high abundance, but low occupancy taxa (i.e. “clumping” (Wright, 1991)) which can be suggestive of niche selection (Morella *et al.*, 2020). Clumping is thought to be impacted by spatial variation in habitat quality, localized reproduction, and stochastic immigration-extinction processes (Wright, 1991). Clumping may also be the result of a competitive lottery-based assembly of the mycobiome (i.e. inhibitory priority effects) which means the first species to arrive will take over the entire niche, excluding other group members (Verster & Borenstein, 2018). In addition to not having the expected abundance-occupancy shape, our data had poor fit to the Sloan neutral model, although the fit was generally consistent with other studies of fungi (Stopnisek & Shade), and also of bacteria (Burns *et al.*, 2016). Thus, the poor fit of the neutral model may indicate that deterministic factors such as competition for niche space, extreme dispersal limitation and variation in habitat quality may be playing a larger role than expected in assembly dynamics of seagrass-associated fungi.

Regardless of model fit, we plotted the global distribution of fourteen core ASVs which were found to be differentially abundant across *Z. marina* tissues with seven deviating from the neutral model (Table 1). Some of these ASVs were found to be globally distributed, while others showed site specificity. For example, both ITS_SV260 (*Saccharomyces paradoxus*) and 18S_SV928 (*Saccharomyces* sp.) were globally distributed and more abundant on leaves and roots (*p* < 0.001, Figure 6, Figure S14). ITS_SV260 was predicted to be neutrally selected while 18S_SV928 was predicted to be plant-selected. *S. paradoxus* is a wild yeast, the sister species to *S. cerevisiae* and has been previously observed as a plant endophyte (Glushakova *et al.*, 2007; Ricks & Koide, 2019). Given the relative abundance of *Saccharomyces* in both the ITS and 18S rRNA gene datasets on *Z. marina* leaves and roots, the global distribution of this taxon and its deviation from the neutral model in the 18S rRNA gene dataset (e.g. 18S_SV928), *Saccharomyces* sp. seems like a good candidate for future work in this system.

In comparison, ITS_SV362 (*Lobulomyces* sp.) was only found at one site, dispersal-limited and is more abundant on leaves (*p* < 0.001, Figure 7). Lobulomycetales have previously been observed in high abundance on and inside *Z. marina* leaves (Ettinger & Eisen, 2019). Members of the Lobulomycetales and marine fungi more generally have been observed to have seasonal dynamics which has may relate to host-dynamics and environmental conditions (Longcore, 1992; Seto & Degawa, 2015; Hassett & Gradinger, 2016; Rojas-Jimenez *et al.*, 2019). Additionally, marine chytrids are known to parasitize seasonal blooms of diatoms (Hassett & Gradinger, 2016; Taylor & Cunliffe, 2016) and diatoms are the dominant eukaryotes observed on seagrass leaf tissues here (Figure S10). Future studies should attempt to confirm whether these and other chytrids assigned to the core mycobiome of *Z. marina* are parasitizing closely associated diatoms or associated with seagrass leaf tissues directly.

In our previous work, we found a *Colletotrichum* sp. ASV to be an abundant endophyte on and in leaves (Ettinger & Eisen, 2019) and postulated that this taxa may be a *Z. marina* specialist (Ettinger & Eisen, 2020). However, no ASVs taxonomically assigned as *Colletotrichum* sp. were defined as part of the global core microbiome, although one ASV, ITS_SV219, was found to deviate from the neutral model and was predicted to be dispersal-limited. Its global distribution supports a pattern of endemism to only a few locations including Bodega Bay, CA, the location of our previous studies (Figure S14). Local adaptation of marine fungi is consistent with patterns of endemism seen in terrestrial fungal studies (Meiser *et al.*, 2014; Grantham *et al.*, 2015), and *Colletotrichum* sp. has been seen before as an endemic endophyte in *Arabidopsis thaliana* (Hiruma *et al.*, 2016). One limitation of ‘core’ community analyses generally, is that it often underplays the importance of rare microbes which can also be essential for host function (Jousset *et al.*, 2017). Thus, future work should include studies of the functional importance of *Colletotrichum* sp. and other rare members of the *Z. marina* mycobiome.

Fungi are not the only eukaryotic microbes associated with *Z*. *marina* and there are many other understudied microorganisms that likely have important roles in the seagrass ecosystem. In fact, seagrass-associated fungi were represented by a mean relative abundance of less than one percent in the 18S rRNA amplicon dataset. This is generally consistent with the proportion of fungi in other marine eukaryotic studies (e.g. 1.3% fungal sequences in Hassett et al. (2020)). Instead, the *Z. marina* associated eukaryotic community was generally dominated by diatoms, oomycetes, and dinoflagellates. Diatom dominance was previously observed in a culture-independent effort of *Z. marina* which found that the bacterial and eukaryotic epibiont communities were highly correlated (Bengtsson *et al.*, 2017). Additionally, oomycetes have been previously cultured in association with *Z*. *marina* and are thought to function as opportunistic pathogens or saprotrophs in this system (Man in ’t Veld *et al.*, 2011, 2019; Ettinger & Eisen, 2020).

Finally, we used FUNGuild to gain insight into possible functional roles of mycobiome, but unfortunately FUNGuild was only able to predict the function of a small portion of ASVs with high confidence. For what could be predicted, the seagrass mycobiome was found to be made of a community of wood saprotrophs, ectomycorrhizal fungi, endophytic fungi and plant pathogens. This functional distribution fits with what might be expected for a plant-associated fungal community, as well as, with what is known of the functional guilds of close relatives of fungal isolates previously isolated from *Z. marina* (Ettinger & Eisen, 2020). However, many dominant members of the fungal community associated with *Z. marina* were not able to be assigned a functional guild, leaving a lot of functional uncertainty to still be explored in this system. Additional studies characterizing seagrass-associated fungi are needed to understand the taxonomic diversity and functional roles of these fungi in the marine ecosystem generally and in particular when associated with seagrasses.

## Author Contributions

Cassandra L. Ettinger analyzed the data, prepared figures and/or tables, performed statistics, wrote and reviewed drafts of the paper.

Laura E. Vann organized experimental design, sample collection and processing, edited and reviewed drafts of the paper.

Jonathan A. Eisen advised on data analysis, edited and reviewed drafts of the paper.

## DNA Deposition

The raw sequences of the ITS2 region and 18S rRNA gene amplicons were deposited at GenBank under accession no. <u>PRJNA667465</u> and <u>PRJNA667462</u> respectively. Sequence reads are also available from the JGI Genome Portal (https://genome.jgi.doe.gov/portal/Popandseaspecies/Popandseaspecies.info.html).

## Acknowledgements

We would like to thank the *Zostera* Experimental Network for helping with sample collection. We are also grateful to Alana J. Firl for assistance in constructing sampling kits, as well as Mikayla Mager and Shuiquan Tang from Zymo Research, Inc. for organizing the DNA extractions. We are very appreciative to Susannah Tringe (ORCID: <u>0000-0001-6479-8427</u>) from the U.S. Department of Energy Joint Genome Institute for helping organize the sequencing component of this project. Guillaume Jospin (ORCID: <u>0000-0002-8746-2632</u>) for his assistance downloading the data files from the JGI web server. John J. Stachowicz and Jeanine L. Olsen for helpful comments and suggestions on this manuscript.

## References

Abarenkov K, Zirk A, Piirmann T, Pöhönen R, Ivanov F, Nilsson RH, Kõljalg U. 2020. UNITE general FASTA release for eukaryotes 2.

Ainsworth TD, Krause L, Bridge T, Torda G, Raina J-B, Zakrzewski M, Gates RD, Padilla-Gamiño JL, Spalding HL, Smith C, et al. 2015. The coral core microbiome identifies rare bacterial taxa as ubiquitous endosymbionts. The ISME journal 9: 2261–2274.

Aitchison J, Barceló-Vidal C, Martín-Fernández JA, Pawlowsky-Glahn V. 2000. Logratio Analysis and Compositional Distance. Mathematical geology 32: 271–275.

Allaire JJ, Xie Y, McPherson J, Luraschi J, Ushey K, Atkins A, Wickham H, Cheng J, Chang W, Iannone R. 2020. rmarkdown: Dynamic Documents for R.

Amend A. 2014. From dandruff to deep-sea vents: Malassezia-like fungi are ecologically hyper-diverse. PLoS pathogens 10: e1004277.

Amend A, Burgaud G, Cunliffe M, Edgcomb VP, Ettinger CL, Gutiérrez MH, Heitman J, Hom EFY, Ianiri G, Jones AC, et al. 2019. Fungi in the Marine Environment: Open Questions and Unsolved Problems. mBio 10.

Anderson MJ. 2001. A new method for non-parametric multivariate analysis of variance. Austral Ecology 26: 32–46.

Bálint M, Tiffin P, Hallström B, O’Hara RB, Olson MS, Fankhauser JD, Piepenbring M, Schmitt I. 2013. Host Genotype Shapes the Foliar Fungal Microbiome of Balsam Poplar (Populus balsamifera). PLoS ONE 8: e53987.

Baselga A, Orme D, Villeger S, De Bortoli J, Leprieur F. 2018. betapart: Partitioning Beta Diversity into Turnover and Nestedness Components.

Bass AJ, Robinson DG, Lianoglou S, Nelson E, Storey JD, from Laurent Gatto WC. 2020. biobroom: Turn Bioconductor objects into tidy data frames.

Becker RA. 2018. maps: Draw Geographical Maps.

Bengtsson MM, Bühler A, Brauer A, Dahlke S, Schubert H, Blindow I. 2017. Eelgrass Leaf Surface Microbiomes Are Locally Variable and Highly Correlated with Epibiotic Eukaryotes. Frontiers in microbiology 8: 1312.

Borovec O, Vohník M. 2018. Ontogenetic transition from specialized root hairs to specific root-fungus symbiosis in the dominant Mediterranean seagrass Posidonia oceanica. Scientific reports 8: 10773.

Bray JR, Roger Bray J, Curtis JT. 1957. An Ordination of the Upland Forest Communities of Southern Wisconsin. Ecological Monographs 27: 325–349.

Burns AR, Stephens WZ, Stagaman K, Wong S, Rawls JF, Guillemin K, Bohannan BJ. 2016. Contribution of neutral processes to the assembly of gut microbial communities in the zebrafish over host development. The ISME journal 10: 655–664.

Callahan BJ, McMurdie PJ, Rosen MJ, Han AW, Johnson AJA, Holmes SP. 2016. DADA2: High-resolution sample inference from Illumina amplicon data. Nature Methods 13: 581–583.

Campanella JJ, Bologna PA, Smalley JV, Rosenzweig EB, Smith SM. 2010. Population structure of Zostera marina (eelgrass) on the western Atlantic coast is characterized by poor connectivity and inbreeding. The Journal of heredity 101: 61–70.

Capone DG. 1982. Nitrogen Fixation (Acetylene Reduction) by Rhizosphere Sediments of the Eelgrass Zostera marina. Marine Ecology Progress Series 10: 67–75.

Chave J. 2004. Neutral theory and community ecology. Ecology Letters 7: 241–253.

Chen H. 2018. VennDiagram: Generate High-Resolution Venn and Euler Plots.

Comeau AM, Vincent WF, Bernier L, Lovejoy C. 2016. Novel chytrid lineages dominate fungal sequences in diverse marine and freshwater habitats. Scientific reports 6: 30120.

Cox F, Newsham KK, Bol R, Dungait JAJ, Robinson CH. 2016. Not poles apart: Antarctic soil fungal communities show similarities to those of the distant Arctic. Ecology letters 19: 528–536.

Crump BC, Wojahn JM, Tomas F, Mueller RS. 2018. Metatranscriptomics and Amplicon Sequencing Reveal Mutualisms in Seagrass Microbiomes. Frontiers in microbiology 9: 388.

Cúcio C, Engelen AH, Costa R, Muyzer G. 2016. Rhizosphere Microbiomes of European Seagrasses Are Selected by the Plant, But Are Not Species Specific. Frontiers in Microbiology 7.

Dray S, Dufour A--B. 2007. The ade4 Package: Implementing the Duality Diagram for Ecologists. Journal of Statistical Software 22: 1–20.

Duffy JE, Reynolds PL, Boström C, Coyer JA, Cusson M, Donadi S, Douglass JG, Eklöf JS, Engelen AH, Eriksson BK, et al. 2015. Biodiversity mediates top-down control in eelgrass ecosystems: a global comparative-experimental approach. Ecology letters 18: 696–705.

Eddelbuettel D. 2013. Seamless R and C++ Integration with Rcpp.

Elzhov TV, Mullen KM, Spiess A-N, Bolker B. 2016. minpack.lm: R Interface to the Levenberg-Marquardt Nonlinear Least-Squares Algorithm Found in MINPACK, Plus Support for Bounds.

Ettinger C. 2020. casett/Global_ZM_fungi_amplicons.

Ettinger CL, Eisen JA. 2019. Characterization of the Mycobiome of the Seagrass,, Reveals Putative Associations With Marine Chytrids. Frontiers in microbiology 10: 2476.

Ettinger CL, Eisen JA. 2020. Fungi, bacteria and oomycota opportunistically isolated from the seagrass, Zostera marina. PloS one 15: e0236135.

Ettinger CL, Voerman SE, Lang JM, Stachowicz JJ, Eisen JA. 2017a. Microbial communities in sediment from patches, but not the leaf or root microbiomes, vary in relation to distance from patch edge. PeerJ 5: e3246.

Ettinger CL, Williams SL, Abbott JM, Stachowicz JJ, Eisen JA. 2017b. Microbiome succession during ammonification in eelgrass bed sediments. PeerJ 5: e3674.

Fahimipour AK, Kardish MR, Lang JM, Green JL, Eisen JA, Stachowicz JJ. 2017. Global-Scale Structure of the Eelgrass Microbiome. Applied and environmental microbiology 83.

Fenchel T, Finlay BJ. 2004. The Ubiquity of Small Species: Patterns of Local and Global Diversity. BioScience 54: 777.

Finlay BJ. 2002. Global dispersal of free-living microbial eukaryote species. Science 296: 1061–1063.

Fourqurean JW, Duarte CM, Kennedy H, Marbà N, Holmer M, Mateo MA, Apostolaki ET, Kendrick GA, Krause-Jensen D, McGlathery KJ, et al. 2012. Seagrass ecosystems as a globally significant carbon stock. Nature Geoscience 5: 505–509.

Frau A, Kenny JG, Lenzi L, Campbell BJ, Ijaz UZ, Duckworth CA, Burkitt MD, Hall N, Anson J, Darby AC, et al. 2019. DNA extraction and amplicon production strategies deeply inf luence the outcome of gut mycobiome studies. Scientific reports 9: 9328.

Gao Z, Li B, Zheng C, Wang G. 2008. Molecular detection of fungal communities in the Hawaiian marine sponges Suberites zeteki and Mycale armata. Applied and environmental microbiology 74: 6091–6101.

Garnier S. 2018. viridis: Default Color Maps from ‘matplotlib’.

Gloor GB, Macklaim JM, Pawlowsky-Glahn V, Egozcue JJ. 2017. Microbiome Datasets Are Compositional: And This Is Not Optional. Frontiers in Microbiology 8.

Glushakova AM, Ivannikova IV, Naumova ES, Chernov II, Naumov GI. 2007. [Massive isolation and identification of Saccharomyces paradoxus yeasts from plant phyllosphere]. Mikrobiologiia 76: 236–242.

Gnavi G, Garzoli L, Poli A, Prigione V, Burgaud G, Varese GC. 2017. The culturable mycobiota of Flabellia petiolata: First survey of marine fungi associated to a Mediterranean green alga. PloS one 12: e0175941.

Goslee SC, Urban DL. 2007. The ecodist package for dissimilarity-based analysis of ecological data. Journal of Statistical Software 22: 1–19.

Grantham NS, Reich BJ, Pacifici K, Laber EB, Menninger HL, Henley JB, Barberán A, Leff JW, Fierer N, Dunn RR. 2015. Fungi identify the geographic origin of dust samples. PloS one 10: e0122605.

Grossart H-P, Van den Wyngaert S, Kagami M, Wurzbacher C, Cunliffe M, Rojas-Jimenez K. 2019. Fungi in aquatic ecosystems. Nature reviews. Microbiology 17: 339–354.

Grossart H-P, Wurzbacher C, James TY, Kagami M. 2016. Discovery of dark matter fungi in aquatic ecosystems demands a reappraisal of the phylogeny and ecology of zoosporic fungi. Fungal Ecology 19: 28–38.

Gumiere T, Durrer A, Bohannan BJM, Andreote FD. 2016. Biogeographical patterns in fungal communities from soils cultivated with sugarcane. Journal of Biogeography 43: 2016–2026.

Gutiérrez MH, Pantoja S, Tejos E, Quiñones RA. 2011. The role of fungi in processing marine organic matter in the upwelling ecosystem off Chile. Marine Biology 158: 205–219.

Harrell FE Jr, from Charles Dupont WC, others. M. 2020. Hmisc: Harrell Miscellaneous.

Hassett BT, Ducluzeau A-LL, Collins RE, Gradinger R. 2017. Spatial distribution of aquatic marine fungi across the western Arctic and sub-arctic. Environmental microbiology 19: 475–484.

Hassett BT, Gradinger R. 2016. Chytrids dominate arctic marine fungal communities. Environmental microbiology 18: 2001–2009.

Hassett BT, Vonnahme TR, Peng X, Gareth Jones EB, Heuzé C. 2020. Global diversity and geography of planktonic marine fungi. Botanica Marina 63: 121–139.

Hawksworth DL, Lücking R. 2017. Fungal Diversity Revisited: 2.2 to 3.8 Million Species. Microbiology spectrum 5.

Hemminga MA, Duarte CM. 2000. Seagrass Ecology. Cambridge University Press.

Hijmans RJ. 2019. geosphere: Spherical Trigonometry.

Hiruma K, Gerlach N, Sacristán S, Nakano RT, Hacquard S, Kracher B, Neumann U, Ramírez D, Bucher M, O’Connell RJ, et al. 2016. Root Endophyte Colletotrichum tofieldiae Confers Plant Fitness Benefits that Are Phosphate Status Dependent. Cell 165: 464–474.

Hothorn T, Hornik K, van de Wiel MA, Zeileis A. 2006. A Lego system for conditional inference. The American Statistician 60: 257–263.

Hothorn T, Hornik K, van de Wiel MA, Zeileis A. 2008. Implementing a class of permutation tests: The coin package. Journal of Statistical Software 28: 1–23.

Huber W, Carey VJ, Gentleman R, Anders S, Carlson M, Carvalho BS, Bravo HC, Davis S, Gatto L, Girke T, et al. 2015. Orchestrating high-throughput genomic analysis with Bioconductor. Nature Methods 12: 115–121.

Hunter PJ, Pink DAC, Bending GD. 2015. Cultivar-level genotype differences influence diversity and composition of lettuce (Lactuca sp.) phyllosphere fungal communities. Fungal Ecology 17: 183–186.

Hurtado-McCormick V, Kahlke T, Petrou K, Jeffries T, Ralph PJ, Seymour JR. 2019. Regional and Microenvironmental Scale Characterization of the Seagrass Microbiome. Frontiers in microbiology 10: 1011.

Huse SM, Ye Y, Zhou Y, Fodor AA. 2012. A core human microbiome as viewed through 16S rRNA sequence clusters. PloS one 7: e34242.

Hyde KD, Gareth Jones EB, Leaño E, Pointing SB, Poonyth AD, Vrijmoed LLP. 1998. Role of fungi in marine ecosystems. Biodiversity and Conservation 7: 1147–1161.

Jones EBG. 2011. Are there more marine fungi to be described? Botanica marina.

Jones EBG, Gareth Jones EB, Pang K-L, Abdel-Wahab MA, Scholz B, Hyde KD, Boekhout T, Ebel R, Rateb ME, Henderson L, et al. 2019. An online resource for marine fungi. Fungal Diversity 96: 347–433.

Jones EBG, Gareth Jones EB, Suetrong S, Sakayaroj J, Bahkali AH, Abdel-Wahab MA, Boekhout T, Pang K-L. 2015. Classification of marine Ascomycota, Basidiomycota, Blastocladiomycota and Chytridiomycota. Fungal Diversity 73: 1–72.

Jousset A, Bienhold C, Chatzinotas A, Gallien L, Gobet A, Kurm V, Küsel K, Rillig MC, Rivett DW, Salles JF, et al. 2017. Where less may be more: how the rare biosphere pulls ecosystems strings. The ISME journal 11: 853–862.

Kagami M, de Bruin A, Ibelings BW, Van Donk E. 2007. Parasitic chytrids: their effects on phytoplankton communities and food-web dynamics. Hydrobiologia 578: 113–129.

Kirichuk NN, Pivkin MV. 2015. Filamentous fungi associated with the seagrass Zostera marina Linnaeus, 1753 of Rifovaya Bay (Peter the Great Bay, the Sea of Japan). Russian Journal of Marine Biology 41: 351–355.

Lahti L, Shetty S. microbiome R package.

Lawrence M, Huber W, Pagès H, Aboyoun P, Carlson M, Gentleman R, Morgan M, Carey V. 2013. Software for Computing and Annotating Genomic Ranges. PLoS Computational Biology 9.

Legendre P, Gallagher ED. 2001. Ecologically meaningful transformations for ordination of species data. Oecologia 129: 271–280.

Littman R, Willis BL, Bourne DG. 2011. Metagenomic analysis of the coral holobiont during a natural bleaching event on the Great Barrier Reef. Environmental Microbiology Reports 3: 651–660.

Longcore JE. 1992. Morphology and Zoospore Ultrastructure of Chytriomyces angularis sp. nov. (Chytridiales). Mycologia 84: 442.

Love MI, Huber W, Anders S. 2014. Moderated estimation of fold change and dispersion for RNA-seq data with DESeq2. Genome Biology 15: 550.

Lowe WH, McPeek MA. 2014. Is dispersal neutral? Trends in Ecology & Evolution 29: 444–450.

Man in ’t Veld WA, Man in ’t W, Karin C H, Brouwer H, de Cock AWAM. 2011. Phytophthora gemini sp. nov., a new species isolated from the halophilic plant Zostera marina in the Netherlands. Fungal Biology 115: 724–732.

Man in ’t Veld WA, Man in ’t W, Karin C H, van Rijswick PCJ, Meffert JP, Boer E, Westenberg M, van der Heide T, Govers LL. 2019. Multiple Halophytophthora spp. and Phytophthora spp. including P. gemini, P. inundata and P. chesapeakensis sp. nov. isolated from the seagrass Zostera marina in the Northern hemisphere. European Journal of Plant Pathology 153: 341–357.

Martin M. 2011. Cutadapt removes adapter sequences from high-throughput sequencing reads. EMBnet.journal 17: 10.

Mata JL, Cebrián J. 2013. Fungal endophytes of the seagrasses Halodule wrightii and Thalassia testudinum in the north-central Gulf of Mexico. Botanica Marina 56.

McMurdie PJ, Holmes S. 2013. phyloseq: An R package for reproducible interactive analysis and graphics of microbiome census data. PLoS ONE 8: e61217.

McMurdie PJ, Holmes S. 2014. Waste Not, Want Not: Why Rarefying Microbiome Data Is Inadmissible. PLoS Computational Biology 10: e1003531.

Meiser A, Bálint M, Schmitt I. 2014. Meta-analysis of deep-sequenced fungal communities indicates limited taxon sharing between studies and the presence of biogeographic patterns. The New phytologist 201: 623–635.

Menkis A, Burokienė D, Gaitnieks T, Uotila A, Johannesson H, Rosling A, Finlay RD, Stenlid J, Vasaitis R. 2012. Occurrence and impact of the root-rot biocontrol agent Phlebiopsis gigantea on soil fungal communities in Picea abies forests of northern Europe. FEMS Microbiology Ecology 81: 438–445.

Morales SE, Biswas A, Herndl GJ, Baltar F. 2019. Global Structuring of Phylogenetic and Functional Diversity of Pelagic Fungi by Depth and Temperature. Frontiers in Marine Science 6.

Morella NM, Weng FC-H, Joubert PM, Metcalf CJE, Lindow S, Koskella B. 2020. Successive passaging of a plant-associated microbiome reveals robust habitat and host genotype-dependent selection. Proceedings of the National Academy of Sciences of the United States of America 117: 1148–1159.

Morgan M, Anders S, Lawrence M, Aboyoun P, Pagès H, Gentleman R. 2009. ShortRead: a Bioconductor package for input, quality assessment and exploration of high-throughput sequence data. Bioinformatics 25: 2607–2608.

Muñiz-Salazar R, Talbot SL, Sage GK, Ward DH, Cabello-Pasini A. 2005. Population genetic structure of annual and perennial populations of Zostera marina L. along the Pacific coast of Baja California and the Gulf of California. Molecular Ecology 14: 711–722.

Nagano Y, Miura T, Nishi S, Lima AO, Nakayama C, Pellizari VH, Fujikura K. 2017. Fungal diversity in deep-sea sediments associated with asphalt seeps at the Sao Paulo Plateau. Deep Sea Research Part II: Topical Studies in Oceanography 146: 59–67.

Neuwirth E. 2014. RColorBrewer: ColorBrewer Palettes.

Nguyen NH, Song Z, Bates ST, Branco S, Tedersoo L, Menke J, Schilling JS, Kennedy PG. 2016. FUNGuild: An open annotation tool for parsing fungal community datasets by ecological guild. Fungal Ecology 20: 241–248.

Ogle DH, Wheeler P, Dinno A. 2020. FSA: Fisheries Stock Analysis.

Oksanen J, Blanchet FG, Friendly M, Kindt R, Legendre P, McGlinn D, Minchin PR, O’Hara RB, Simpson GL, Solymos P, et al. 2019. vegan: Community Ecology Package.

Olsen JL, Stam WT, Coyer JA, Reusch TBH, Billingham M, Boström C, Calvert E, Christie H, Granger S, la Lumière R, et al. 2004. North Atlantic phylogeography and large-scale population differentiation of the seagrass Zostera marina L. Molecular ecology 13: 1923–1941.

Orsi W, Biddle JF, Edgcomb V. 2013. Deep sequencing of subseafloor eukaryotic rRNA reveals active Fungi across marine subsurface provinces. PloS one 8: e56335.

Ort BS, Cohen CS, Boyer KE, Wyllie-Echeverria S. 2012. Population Structure and Genetic Diversity among Eelgrass (Zostera marina) Beds and Depths in San Francisco Bay. Journal of Heredity 103: 533–546.

Orth RJ, Carruthers TJB, Dennison WC, Duarte CM, Fourqurean JW, Heck KL, Randall Hughes A, Kendrick GA, Judson Kenworthy W, Olyarnik S, et al. 2006. A Global Crisis for Seagrass Ecosystems. BioScience 56: 987.

Pauvert C, Buée M, Laval V, Edel-Hermann V, Fauchery L, Gautier A, Lesur I, Vallance J, Vacher C. 2019. Bioinformatics matters: The accuracy of plant and soil fungal community data is highly dependent on the metabarcoding pipeline. Fungal Ecology 41: 23–33.

Peay KG, Bidartondo MI, Arnold AE. 2010. Not every fungus is everywhere: scaling to the biogeography of fungal-plant interactions across roots, shoots and ecosystems. The New phytologist 185: 878–882.

Peay KG, Kennedy PG, Talbot JM. 2016. Dimensions of biodiversity in the Earth mycobiome. Nature reviews. Microbiology 14: 434–447.

Pedersen TL. 2020. patchwork: The Composer of Plots.

Petersen L-E, Marner M, Labes A, Tasdemir D. 2019. Rapid Metabolome and Bioactivity Profiling of Fungi Associated with the Leaf and Rhizosphere of the Baltic Seagrass Zostera marina. Marine Drugs 17: 419.

Picard KT. 2017. Coastal marine habitats harbor novel early-diverging fungal diversity. Fungal Ecology 25: 1–13.

Quast C, Pruesse E, Yilmaz P, Gerken J, Schweer T, Yarza P, Peplies J, Glöckner FO. 2013. The SILVA ribosomal RNA gene database project: improved data processing and web-based tools. Nucleic acids research 41: D590–6.

Raghukumar S. 2017. The Marine Environment and the Role of Fungi. Fungi in Coastal and Oceanic Marine Ecosystems: 17–38.

Rao CR. 1997. An Alternative to Correspondence Analysis Using Hellinger Distance.

Ricks KD, Koide RT. 2019. The role of inoculum dispersal and plant species identity in the assembly of leaf endophytic fungal communities. PloS one 14: e0219832.

Risely A. 2020. Applying the core microbiome to understand host–microbe systems. Journal of Animal Ecology 89: 1549–1558.

Ritchie ME, Phipson B, Wu D, Hu Y, Law CW, Shi W, Smyth GK. 2015. limma powers differential expression analyses for RNA-sequencing and microarray studies. Nucleic Acids Research 43: e47.

Robinson D, Hayes A. 2020. broom: Convert Statistical Analysis Objects into Tidy Tibbles.

Rojas-Jimenez K, Rieck A, Wurzbacher C, Jürgens K, Labrenz M, Grossart H-P. 2019. A Salinity Threshold Separating Fungal Communities in the Baltic Sea. Frontiers in microbiology 10: 680.

Rosindell J, Hubbell SP, Etienne RS. 2011. The unified neutral theory of biodiversity and biogeography at age ten. Trends in ecology & evolution 26: 340–348.

Sakayaroj J, Preedanon S, Supaphon O, Jones EBG, Phongpaichit S. 2010. Phylogenetic diversity of endophyte assemblages associated with the tropical seagrass Enhalus acoroides in Thailand. Fungal Diversity 42: 27–45.

Salazar G. 2020. EcolUtils: Utilities for community ecology analysis.

Sapkota R, Knorr K, Jørgensen LN, O’Hanlon KA, Nicolaisen M. 2015. Host genotype is an important determinant of the cereal phyllosphere mycobiome. The New phytologist 207: 1134–1144.

Sarkar D. 2008. Lattice: Multivariate Data Visualization with R.

Schmidt VT, Smith KF, Melvin DW, Amaral-Zettler LA. 2015. Community assembly of a euryhaline fish microbiome during salinity acclimation. Molecular ecology 24: 2537–2550.

Seto K, Degawa Y. 2015. Cyclopsomyces plurioperculatus: a new genus and species of Lobulomycetales (Chytridiomycota, Chytridiomycetes) from Japan. Mycologia 107: 633–640.

Shade A, Stopnisek N. 2019. Abundance-occupancy distributions to prioritize plant core microbiome membership. Current opinion in microbiology 49: 50–58.

Shoemaker G, Wyllie-Echeverria S. 2013. Occurrence of rhizomal endophytes in three temperate northeast pacific seagrasses. Aquatic Botany 111: 71–73.

Simpson GL. 2019. permute: Functions for Generating Restricted Permutations of Data.

Sloan WT, Woodcock S, Lunn M, Head IM, Curtis TP. 2007. Modeling taxa-abundance distributions in microbial communities using environmental sequence data. Microbial ecology 53: 443–455.

Sprockett D. 2020. reltools: Microbiome Amplicon Analysis and Visualization.

Stoeck T, Bass D, Nebel M, Christen R, Jones MDM, Breiner H-W, Richards TA. 2010. Multiple marker parallel tag environmental DNA sequencing reveals a highly complex eukaryotic community in marine anoxic water. Molecular ecology 19 Suppl 1: 21–31.

Stopnisek N, Shade A. Prioritizing persistent microbiome members in the common bean rhizosphere: an integrated analysis of space, time, and plant genotype.

Sun F, Zhang X, Zhang Q, Liu F, Zhang J, Gong J. 2015. Seagrass (Zostera marina) Colonization Promotes the Accumulation of Diazotrophic Bacteria and Alters the Relative Abundances of Specific Bacterial Lineages Involved in Benthic Carbon and Sulfur Cycling. Applied and environmental microbiology 81: 6901–6914.

Supaphon P, Phongpaichit S, Sakayaroj J, Rukachaisirikul V, Kobmoo N, Spatafora JW. 2017. Phylogenetic community structure of fungal endophytes in seagrass species. Botanica Marina 60.

Talbot JM, Bruns TD, Taylor JW, Smith DP, Branco S, Glassman SI, Erlandson S, Vilgalys R, Liao H-L, Smith ME, et al. 2014. Endemism and functional convergence across the North American soil mycobiome. Proceedings of the National Academy of Sciences of the United States of America 111: 6341–6346.

Taylor JD, Cunliffe M. 2016. Multi-year assessment of coastal planktonic fungi reveals environmental drivers of diversity and abundance. The ISME journal 10: 2118–2128.

Tedersoo L, Bahram M, Põlme S, Kõljalg U, Yorou NS, Wijesundera R, Villarreal Ruiz L, Vasco-Palacios AM, Thu PQ, Suija A, et al. 2014. Fungal biogeography. Global diversity and geography of soil fungi. Science 346: 1256688.

Therneau TM. 2020. A Package for Survival Analysis in R.

Tisthammer KH, Cobian GM, Amend AS. 2016. Global biogeography of marine fungi is shaped by the environment. Fungal Ecology 19: 39–46.

Trevathan-Tackett SM, Allnutt TR, Sherman CDH, Richardson ME, Crowley TM, Macreadie PI. 2020. Spatial variation of bacterial and fungal communities of estuarine seagrass leaf microbiomes. Aquatic Microbial Ecology 84: 59–74.

Verster AJ, Borenstein E. 2018. Competitive lottery-based assembly of selected clades in the human gut microbiome. Microbiome 6: 186.

Wainwright BJ, Bauman AG, Zahn GL, Todd PA, Huang D. 2019a. Characterization of fungal biodiversity and communities associated with the reef macroalga Sargassum ilicifolium reveals fungal community differentiation according to geographic locality and algal structure. Marine Biodiversity 49: 2601–2608.

Wainwright BJ, Zahn GL, Arlyza IS, Amend AS. 2018. Seagrass-associated fungal communities follow Wallace’s line, but host genotype does not structure fungal community. Journal of Biogeography 45: 762–770.

Wainwright BJ, Zahn GL, Zushi J, Lee NLY, Ooi JLS, Lee JN, Huang D. 2019b. Seagrass-associated fungal communities show distance decay of similarity that has implications for seagrass management and restoration. Ecology and Evolution 9: 11288–11297.

Wang Q, Garrity GM, Tiedje JM, Cole JR. 2007. Naive Bayesian classifier for rapid assignment of rRNA sequences into the new bacterial taxonomy. Applied and environmental microbiology 73: 5261–5267.

Wang L, Tomas F, Mueller RS. 2020. Nutrient enrichment increases size of Zostera marina shoots and enriches for sulfur and nitrogen cycling bacteria in root-associated microbiomes. FEMS microbiology ecology 96.

White TJ, Bruns T, Lee S, Taylor J. 1990. Amplification and direct sequencing of fungal ribosomal RNA genes for phylogenetics. In: PCR protocols: a guide to methods and applications. 315–322.

Wickham H. 2007. Reshaping data with the reshape package. Journal of Statistical Software 21.

Wickham H. 2016. ggplot2: Elegant Graphics for Data Analysis.

Wickham H, Averick M, Bryan J, Chang W, McGowan LD, François R, Grolemund G, Hayes A, Henry L, Hester J, et al. 2019. Welcome to the tidyverse. Journal of Open Source Software 4: 1686.

Wickham H, Seidel D. 2020. scales: Scale Functions for Visualization.

Wright DH. 1991. Correlations Between Incidence and Abundance are Expected by Chance. Journal of Biogeography 18: 463.

Xie Y. 2014. knitr: A Comprehensive Tool for Reproducible Research in R (V Stodden, F Leisch, and RD Peng, Eds.). Implementing Reproducible Computational Research.

Yarden O. 2014. Fungal association with sessile marine invertebrates. Frontiers in microbiology 5: 228.

Yilmaz P, Parfrey LW, Yarza P, Gerken J, Pruesse E, Quast C, Schweer T, Peplies J, Ludwig W, Glöckner FO. 2014. The SILVA and ‘All-species Living Tree Project (LTP)’ taxonomic frameworks. Nucleic Acids Research 42: D643–D648.

Yu G. 2020. ggplotify: Convert Plot to ‘grob’ or ‘ggplot’ Object.

Zeileis A, Croissant Y. 2010. Extended Model Formulas in R: Multiple Parts and Multiple Responses. Journal of Statistical Software 34: 1–13.

Zhou J, Ning D. 2017. Stochastic Community Assembly: Does It Matter in Microbial Ecology? Microbiology and molecular biology reviews: MMBR 81.

Zuccaro A, Schoch CL, Spatafora JW, Kohlmeyer J, Draeger S, Mitchell JI. 2008. Detection and identification of fungi intimately associated with the brown seaweed Fucus serratus. Applied and environmental microbiology 74: 931–941.

