## Supplemental Materials for "Global diversity and biogeography of the *Zostera marina* mycobiome"

**Supplemental Figures:**

**Figure S1.** Within sample diversity varies across sites. Boxplot visualizations of mean Shannon diversities for each sample type (leaf, root, sediment) at each site collected (Table S1) based on (A) ITS2 region amplicon data and (B) 18S rRNA gene amplicon data. The standard error of the mean Shannon diversity at each site for each sample type is represented by error bars, and bars are colored by sample type.

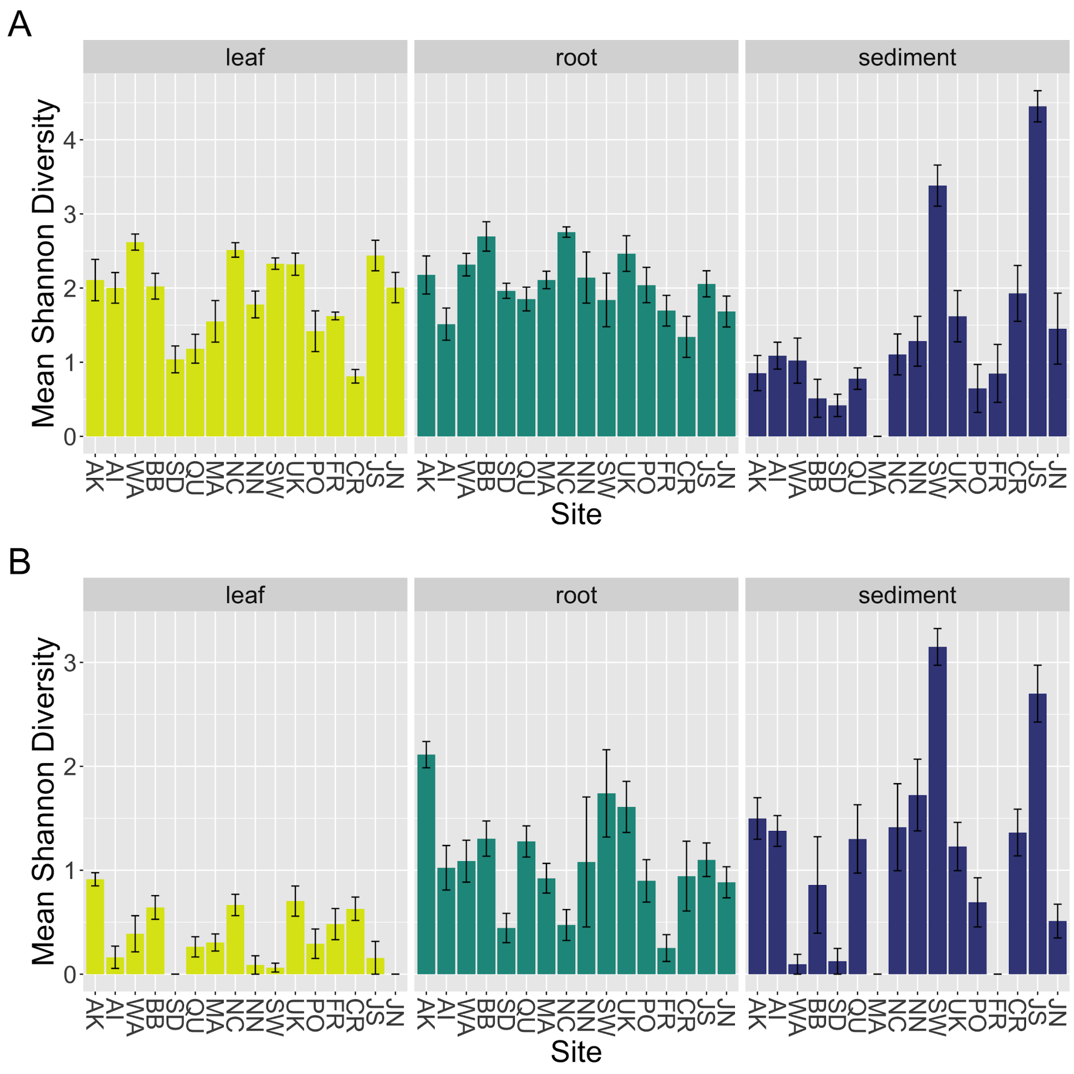

**Figure S2.** Community structure varies between sites and oceans. Principal coordinates analysis (PCoA) visualization of Hellinger distances of fungal communities associated with leaves, roots and sediment. For (A) ITS2 region amplicon data and (B) 18S rRNA amplicon data, points in the ordinations are colored by site collected (Table S1) and represented by shapes based on sample type (circles), root (triangles) or sediment (squares). For (C) ITS2 region amplicon data and (D) 18S rRNA amplicon data, points in the ordinations are colored by ocean (Pacific: blue, Atlantic: orange) and represented by shapes based on sample type (circles), root (triangles) or sediment (squares).

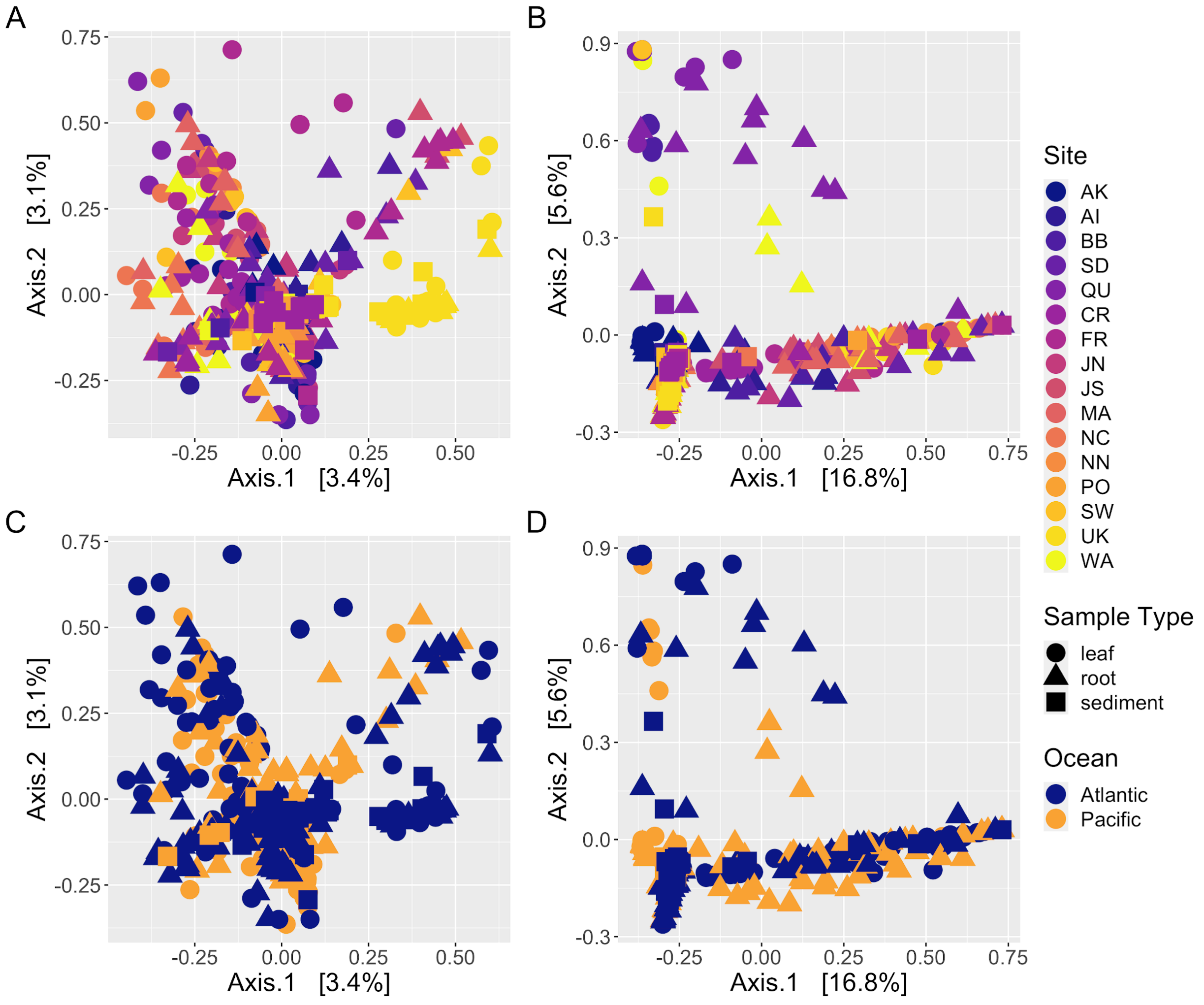

**Figure S3.** Mantel tests suggest a distance-decay relationship. Scatterplots depicting the weak, but significant positive distance–decay relationship between fungal community beta diversity (Hellinger distance) using the 18S rRNA gene amplicon data and geographical distance (km) between sites for leaves from the (A) Pacific Ocean, and (B) Atlantic Ocean, roots from the (C) Pacific Ocean, and (D) Atlantic Ocean, and sediment from the (E) Pacific Ocean and (F) Atlantic Ocean.

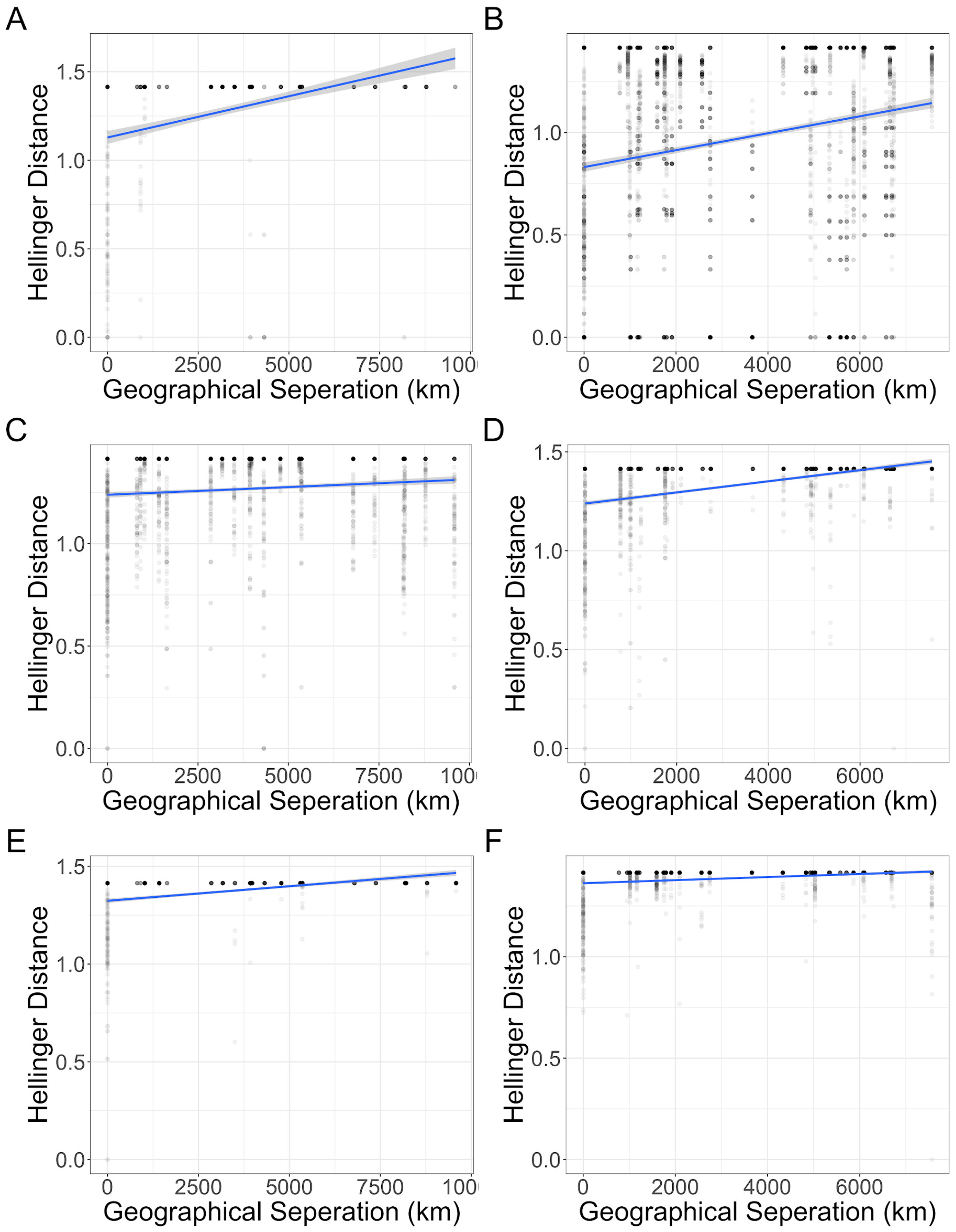

**Figure S4.** Mantel correlograms suggest distance-decay relationships are driven by samples from closest geographic sites. Mantel correlograms for the ITS2 region amplicon data for the (A) Pacific Ocean, and (B) Atlantic Ocean. The x-axis is binned geographic distance class indices and the y-axis is the Mantel correlation statistic. Black points are statistically significant ($p < 0.05$) and white points are not statistically significant.

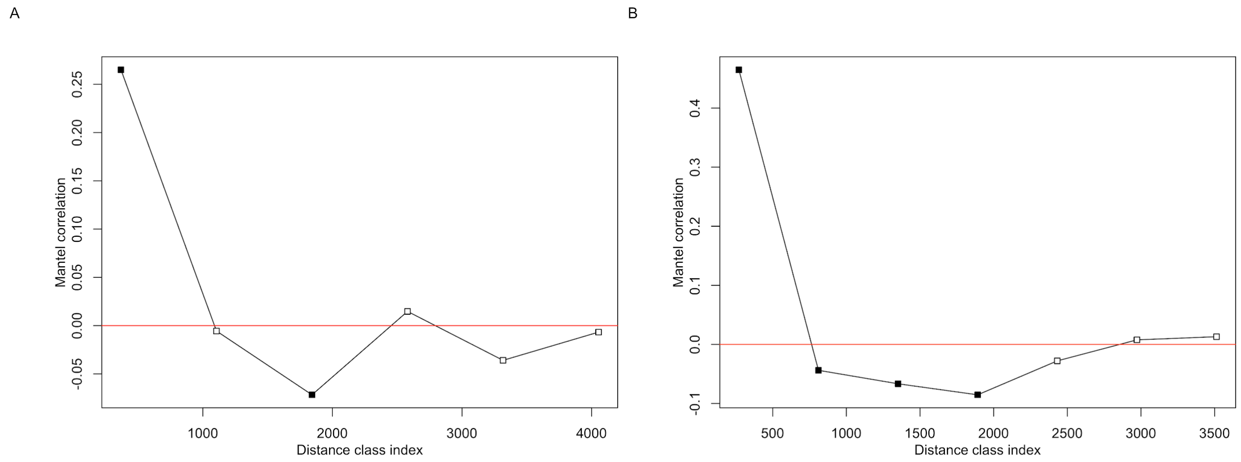

**Figure S5.** Mantel tests suggest a distance-decay relationship. Scatterplots depicting the weak, but significant positive distance–decay relationship between fungal community beta diversity (Hellinger distance) using the ITS2 region amplicon data and geographical distance (km) between sites for roots from the (A) Pacific Ocean, and (B) Atlantic Ocean, and sediment from the (C) Pacific Ocean, and (D) Atlantic Ocean.

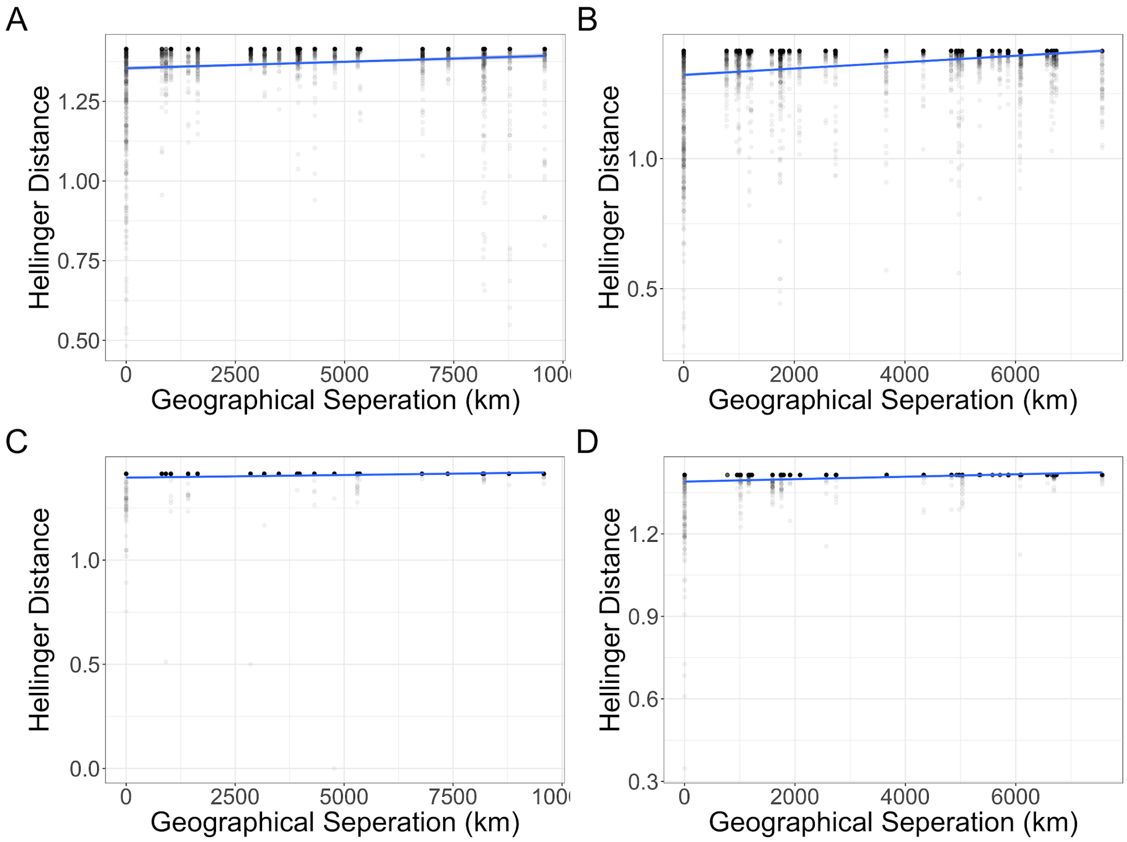

**Figure S6.** Fungal community composition differs between tissue types. The mean relative abundance of taxonomic orders are shown for the (A) ITS2 region amplicon data and (B) the 18S rRNA amplicon data. Only orders with a mean relative abundance of greater than one percent are shown across bulk sample types (leaf, root, and sediment), with the standard error of the mean represented by error bars and bars colored by taxonomic phylum.

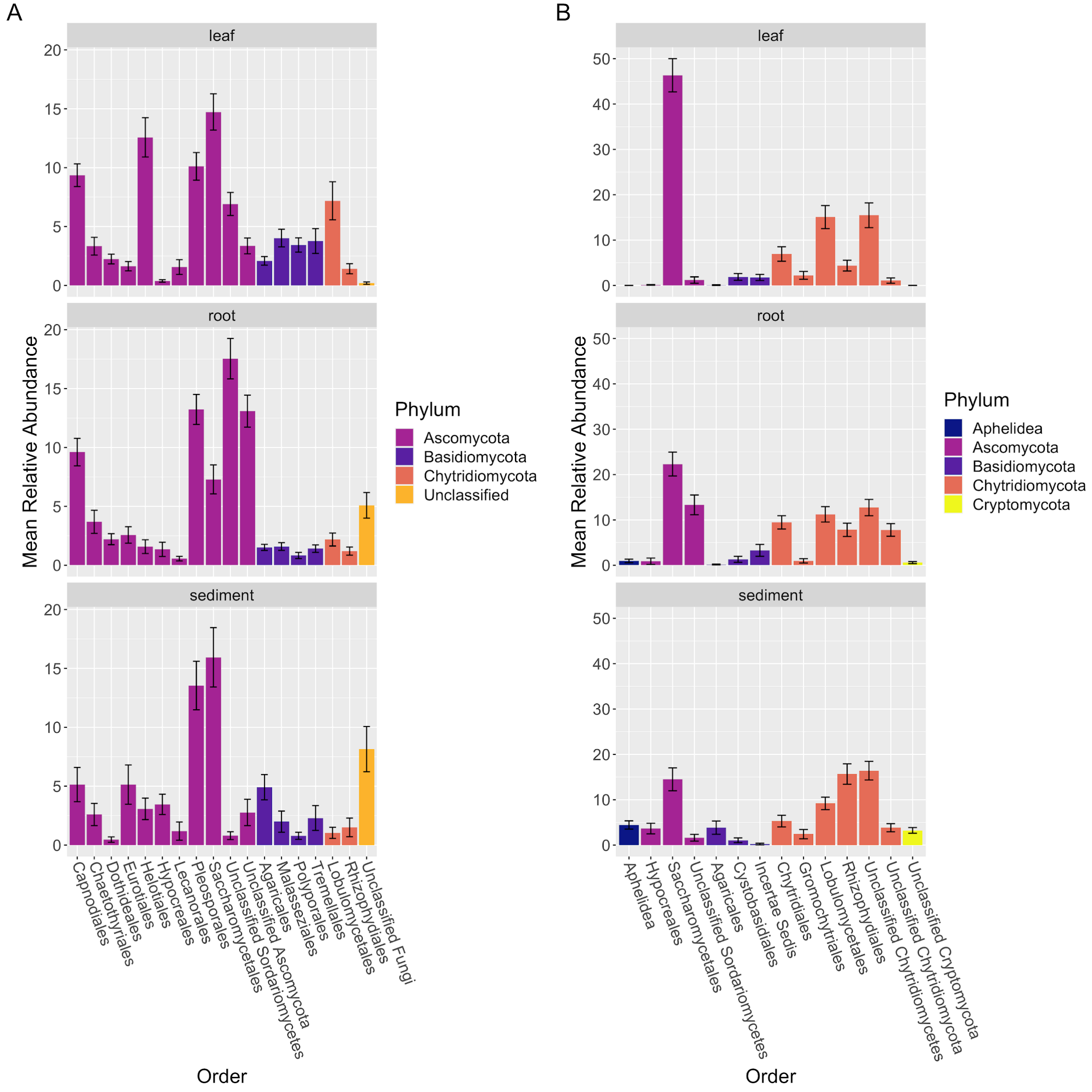

**Figure S7.** Abundance-occupancy distributions reveal core mycobiomes. Abundance-occupancy distributions were used to define core members of the (A) leaf, (B) root and (C) sediment mycobiomes for the 18S rRNA gene amplicon data. Each point represents an ASV with core members indicated by a color (leaf = yellow, root = green, sediment = blue) and non-core ASVs in white. Ranked ASVs were predicted to be in the core based on a final percent increase of equal or greater than 10%. The solid line represents the fit of the neutral model, and the dashed line is 95% confidence around the model prediction. ASVs above the neutral model are predicted to be selected for by the environment (e.g. by the host plant, *Z. marina*), and those below the model are predicted to be selected-against or dispersal-limited.

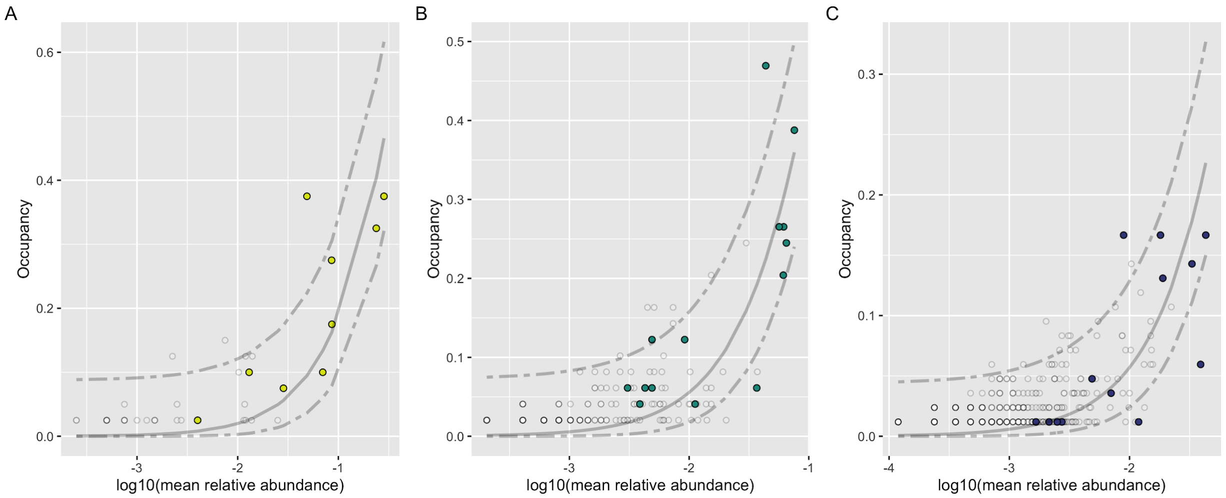

**Figure S8.** Differentially abundant ASVs across tissues. Fungal ITS2 region ASVs were identified using DESeq2 whose abundance differed significantly between pair-wise sample types (leaf, root, sediment). Each plot shows the $\log_2$ fold change of ASVs which were differentially abundant between (A) leaves and rhizosphere sediment, (B) leaves and roots, and (C) roots and rhizosphere sediment. A positive $\log_2$ fold change means the ASV was more abundant in the first tissue and a negative $\log_2$ fold change means the ASV was more abundant in the second tissue in the comparison.

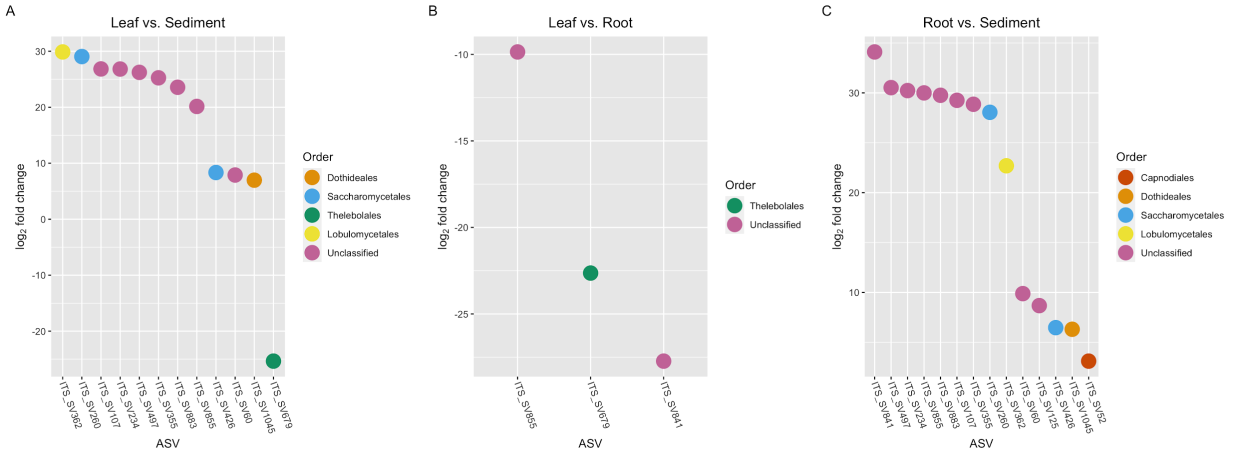

**Figure S9.** Differentially abundant ASVs across tissues. Fungal 18S rRNA gene ASVs were identified  using DESeq2 whose abundance differed significantly between pair-wise sample types (leaf, root, sediment). Each plot shows the $\log_2$ fold change of ASVs which were differentially abundant between (A) leaves and rhizosphere sediment, and (B) roots and rhizosphere sediment. A positive $\log_2$ fold change means the ASV was more abundant in the first tissue and a negative $\log_2$ fold change means the ASV was more abundant in the second tissue in the comparison. No ASVs were differentially abundant between leaves and roots.

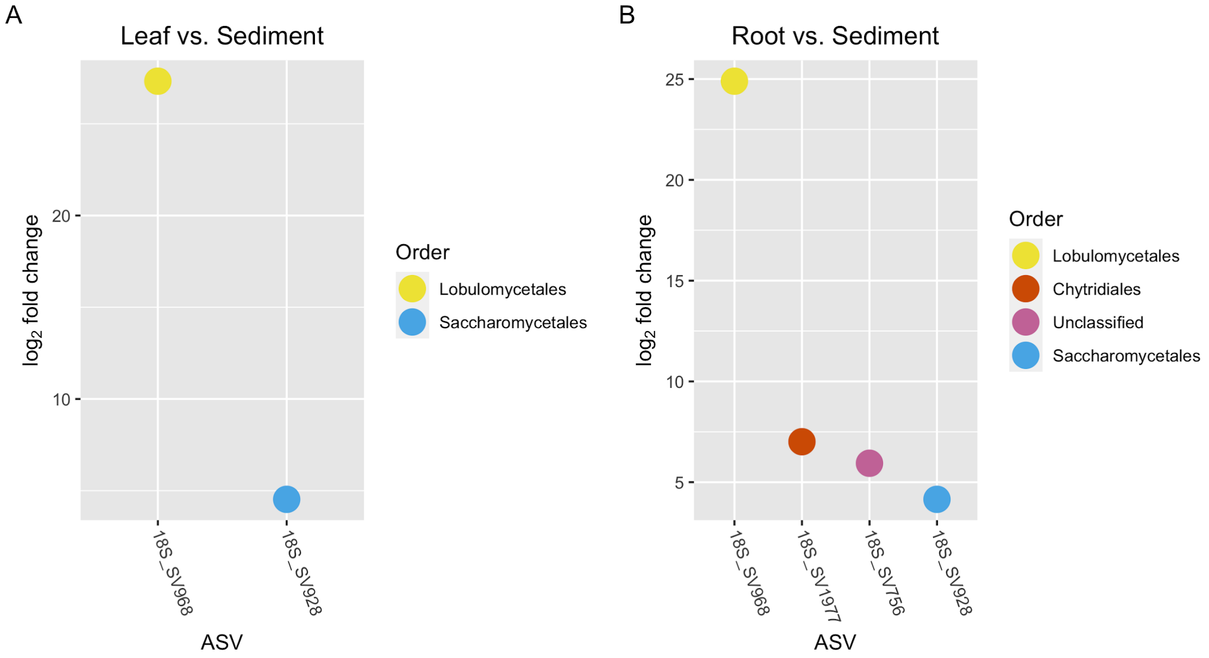

**Figure S10.** Eukaryotic community composition differs between tissues. The mean relative abundance of broad taxonomic groups are shown for the 18S rRNA gene amplicon data. Only groups with a mean relative abundance of greater than 0.1 percent are shown across sample types (leaf, root and sediment), with the standard error of the mean represented by error bars and bars colored by taxonomic groups.

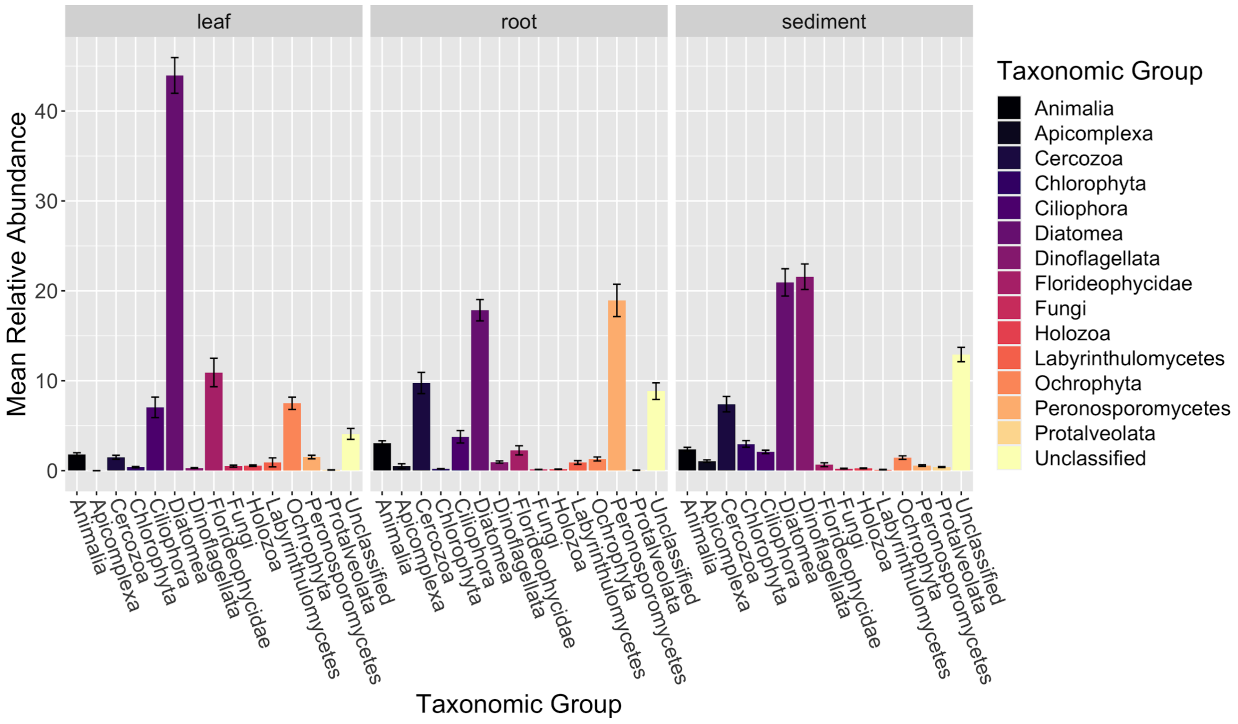

**Figure S11.** Counts of fungal trophic modes based on the ITS2 region amplicon data. The count of fungal trophic modes based on FUNGuild, (A) across fungal guilds assigned with high probability, and (B) across all probability levels for ASVs with a mean relative abundance of 0.1 percent or greater.

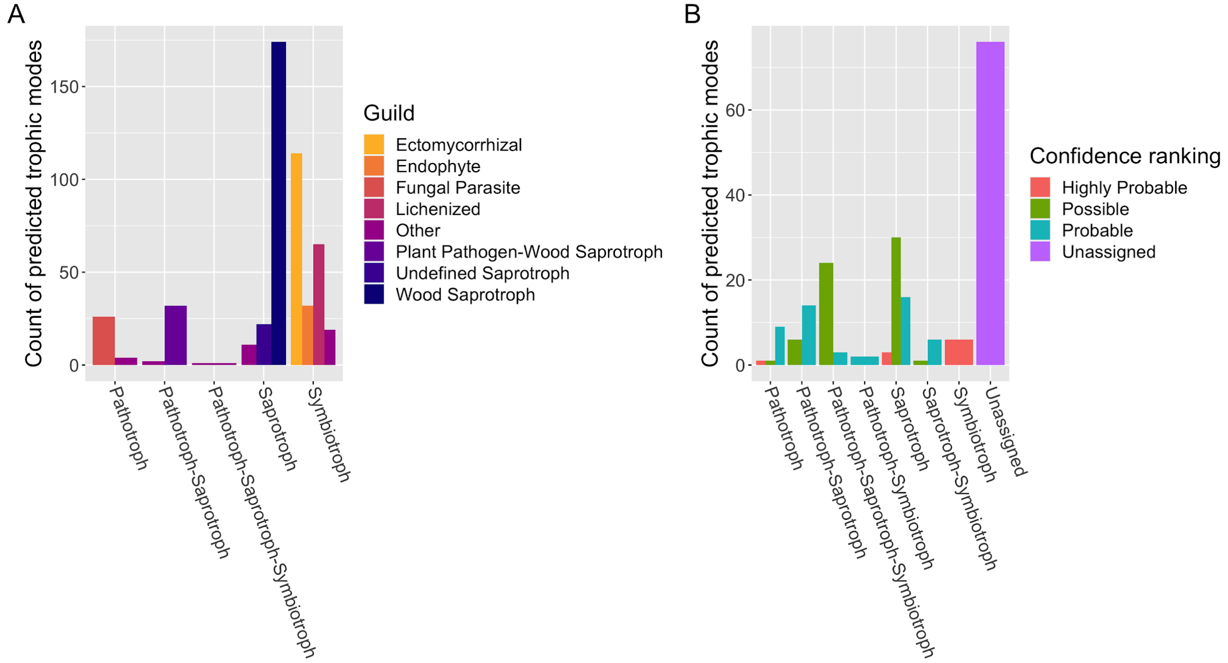

**Figure S12.** Counts of fungal trophic modes based on the 18S rRNA gene amplicon data. The count of fungal trophic modes based on FUNGuild, (A) across fungal guilds assigned with high probability, and (B) across all probability levels for ASVs with a mean relative abundance of 0.1 percent or greater.

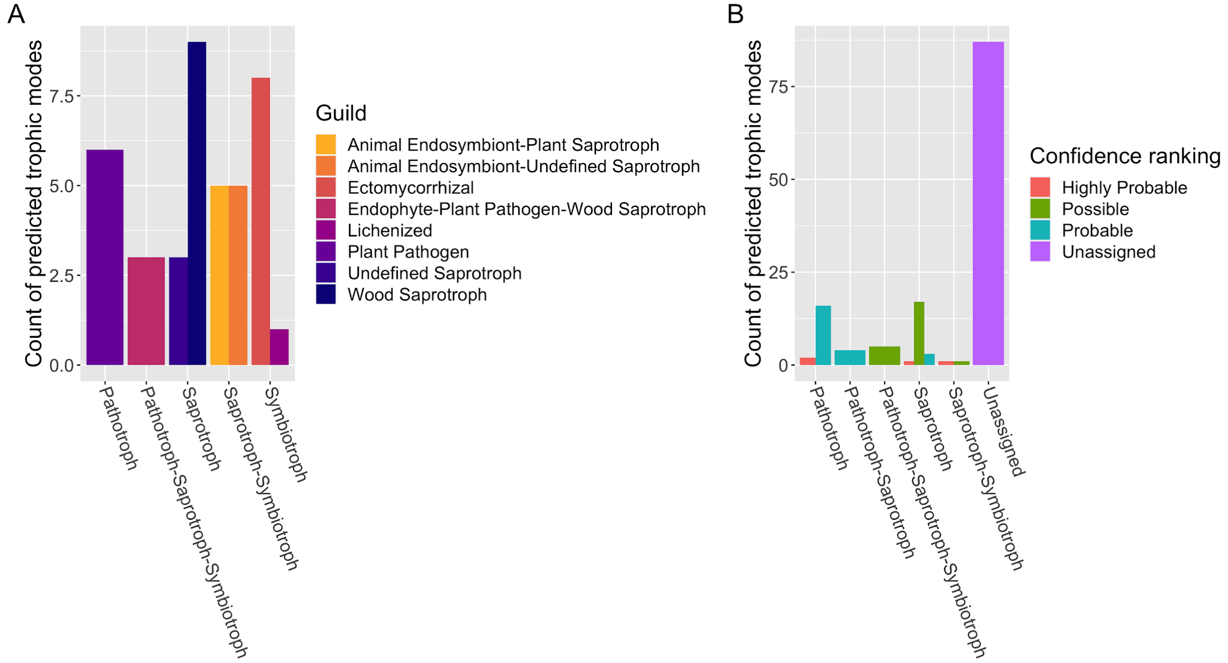

**Figure S13.** Example of differentially abundant plant-selected core ASV. Here we show the global distribution of 18S_SV928, an ASV predicted to be plant-selected and a member of the core mycobiomes of both leaves and roots and also differentially abundant between leaves and sediment ($p < 0.001$), and roots and sediment ($p < 0.001$) using DESeq2. In (A) we plot the mean relative abundance of 18S_SV928 at each site on leaves on a global map, and in (B) we plot the mean relative abundance of 18S_SV928 on leaves, roots and sediment across sites, with the standard error of the mean represented by error bars and bars colored by sample type.

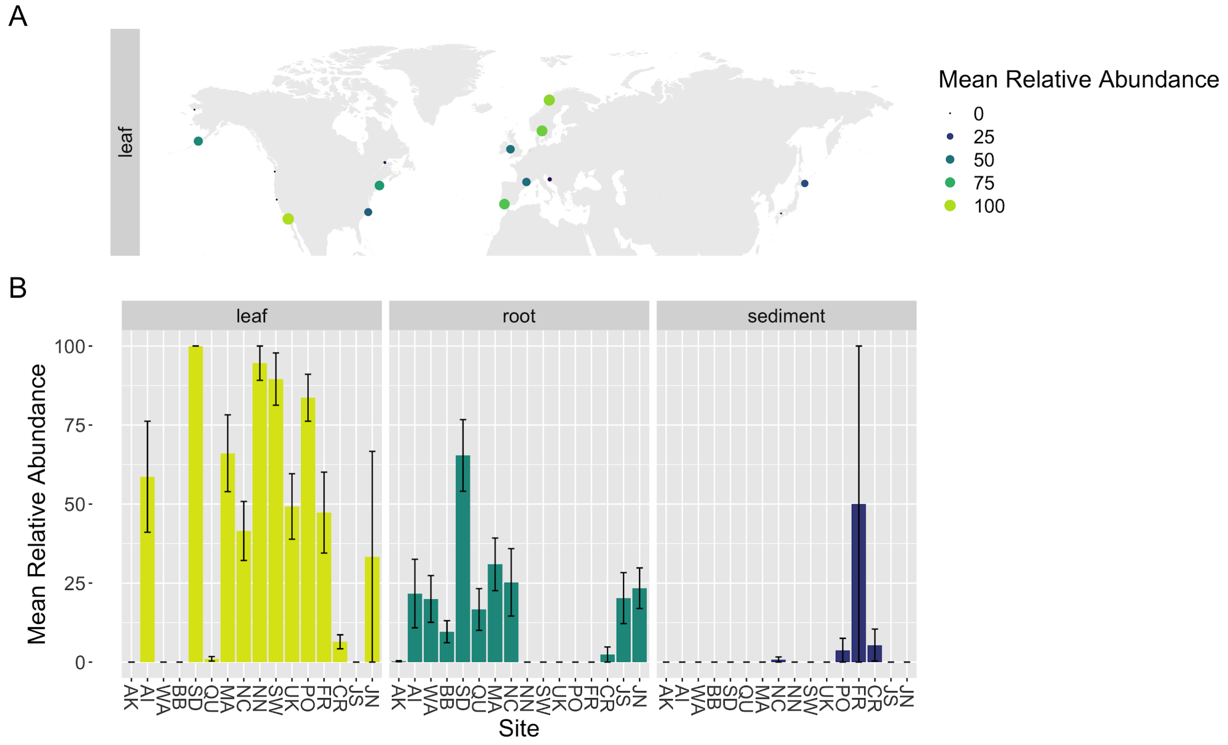

**Figure S14.** Example of dispersal-limited ASV. Here we show the global distribution of ITS_SV219, taxonomically identified as a *Colletotrichum sp*. and predicted to be dispersal-limited. In (A) we plot the mean relative abundance of ITS_SV219 at each site on leaves on a global map, and in (B) we plot the mean relative abundance of ITS_SV219 on leaves, roots and sediment across sites, with the standard error of the mean represented by error bars and bars colored by sample type.

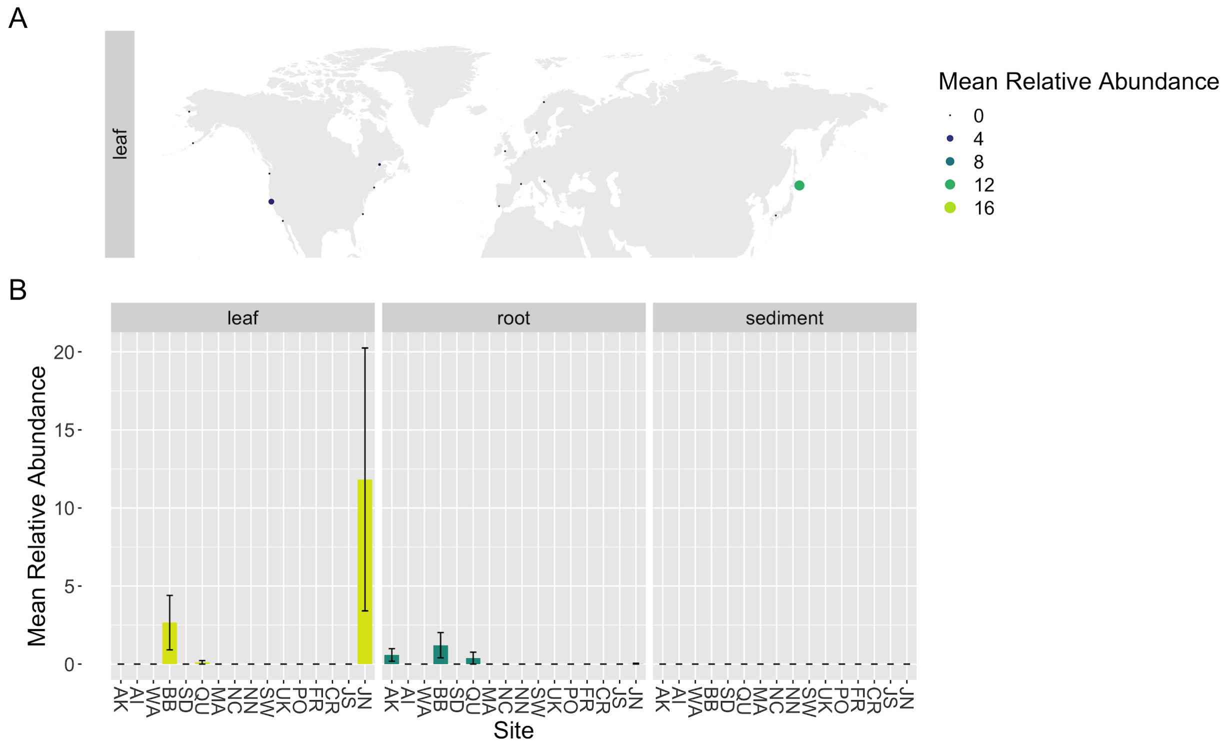

**Supplemental Tables:**

**Table S1.** Description of collection sites. Here we report on the specifics of each collection site including the site code, site name, month of sample collection, year of sample collection, and site latitude and longitude.

| Site Code | Site | Site Name | Collection Month | Collection Year | Latitude | Longitude |
|---|---|---|---|---|---|---|
| **AK** | Alaska - North | Kotzebue | October | 2016 | 64.485428 | -164.76189 |
| **AI** | Alaska - South | Izembek | October | 2016 | 55.328899 | -162.82121 |
| **BB** | California - North | Westside Park, Bodega Bay | December | 2016 | 38.319755 | -123.05514 |
| **SD** | California - South | Shelter Island | November | 2016 | 32.713756 | -117.22547 |
| **QU** | Canada | Pointe-Lebel | September | 2016 | 49.11237 | -68.17593 |
| **CR** | Croatia | Posedarje | September | 2016 | 44.21155 | 15.4906946 |
| **FR** | French Mediterranean | Bouzigues, Etang de Thau | October | 2016 | 43.446971 | 3.661503 |

| | | | | | | |
|---|---|---|---|---|---|---|
| **JN** | Japan - North | Akkeshi-ko estuary | September | 2016 | 43.021167 | 144.903217 |
| **JS** | Japan - South | Ikunoshima | September | 2016 | 34.297834 | 132.91631 |
| **MA** | Massachusetts | Dorothy Cove | October | 2016 | 42.42014 | -70.91544 |
| **NC** | North Carolina | Middle Marsh | April | 2017 | 34.692458 | -76.622589 |
| **NN** | Norway | Ršvika | July | 2016 | 67.2667233 | 15.2560633 |
| **PO** | Portugal | Culatatra | October | 2016 | 37.01427 | -7.493273 |
| **SW** | Sweden | Torseršd | August | 2016 | 58.3131 | 11.5488 |
| **UK** | Wales | Porth Dinllaen | March | 2017 | 52.990731 | -4.450321 |
| **WA** | Washington | Willapa Bay | September | 2016 | 46.474 | -124.028 |

**Table S2.** Evidence of distance-decay from Mantel test results. After first subsetting each dataset (ITS2 region, 18S rRNA gene) by ocean (Pacfic, Atlantic) and sample type (leaf, root, sediment), we identified distance-decay patterns by testing for correlations between the community distance (Bray-Curtis, Hellinger) and the geographic distance between samples. Here we report the results of each Mantel test.

| Ocean | Amplicon dataset | Sample type | Beta diversity metric | Mantel statistic (*r*) | *p*-value |
|---|---|---|---|---|---|
| **Pacific** | ITS2 region | Leaf | Bray-Curtis | 0.197 | 0.0001 |
| **Pacific** | ITS2 region | Leaf | Hellinger | 0.1767 | 0.0001 |
| **Pacific** | 18S rRNA gene | Leaf | Bray-Curtis | 0.1655 | 0.0001 |
| **Pacific** | 18S rRNA gene | Leaf | Hellinger | 0.1326 | 0.0001 |
| **Pacific** | ITS2 region | Root | Bray-Curtis | 0.2195 | 0.0001 |
| **Pacific** | ITS2 region | Root | Hellinger | 0.1869 | 0.0001 |
| **Pacific** | 18S rRNA gene | Root | Bray-Curtis | 0.4589 | 0.0001 |
| **Pacific** | 18S rRNA gene | Root | Hellinger | 0.4202 | 0.0001 |
| **Pacific** | ITS2 region | Sediment | Bray-Curtis | 0.1417 | 0.0002 |
| **Pacific** | ITS2 region | Sediment | Hellinger | -0.03682 | 0.8267 |
| **Pacific** | 18S rRNA gene | Sediment | Bray-Curtis | 0.06958 | 0.0308 |

| | | | | | |
|---|---|---|---|---|---|
| **Pacific** | 18S rRNA gene | Sediment | Hellinger | 0.09003 | 0.0111 |
| **Atlantic** | ITS2 region | Leaf | Bray-Curtis | 0.1139 | 0.0001 |
| **Atlantic** | ITS2 region | Leaf | Hellinger | 0.1057 | 0.0001 |
| **Atlantic** | 18S rRNA gene | Leaf | Bray-Curtis | 0.06604 | 0.0004 |
| **Atlantic** | 18S rRNA gene | Leaf | Hellinger | 0.05843 | 0.0007 |
| **Atlantic** | ITS2 region | Root | Bray-Curtis | 0.1655 | 0.0001 |
| **Atlantic** | ITS2 region | Root | Hellinger | 0.1646 | 0.0001 |
| **Atlantic** | 18S rRNA gene | Root | Bray-Curtis | 0.1639 | 0.0001 |
| **Atlantic** | 18S rRNA gene | Root | Hellinger | 0.1793 | 0.0001 |
| **Atlantic** | ITS2 region | Sediment | Bray-Curtis | 0.2713 | 0.0001 |
| **Atlantic** | ITS2 region | Sediment | Hellinger | 0.2094 | 0.0001 |
| **Atlantic** | 18S rRNA gene | Sediment | Bray-Curtis | 0.4632 | 0.0001 |
| **Atlantic** | 18S rRNA gene | Sediment | Hellinger | 0.4216 | 0.0001 |

**Table S3.** Predicted core ITS region ASVs. ASVs were ranked by their abundance-occupancy distribution and then predicted to be in the core based on a final percent increase of equal or greater than 10% to Bray-Curtis dissimilarity. The Sloan neutral model was then applied to the abundance-occupancy distributions to identify ASVs that deviate such that ASVs above the neutral model are predicted to be selected for by the environment (e.g. by the host plant, *Z. marina*), and those below the model are predicted to be selected-against or dispersal-limited. Here we report the core ASVs for the *Z. marina* leaf, root and sediment mycobiomes, whether these ASVs deviate from the neutral model (above, below, none) and the taxonomy of each ASV.

| ASV | Core prediction | Neutral model deviation | Taxonomy |
|---|---|---|---|
| **ITS_SV45** | leaf, root, sediment | above, above, above | *Cladosporium* sp. |
| **ITS_SV52** | leaf, root, sediment | above, above, none | *Mycosphaerella tassiana* |
| **ITS_SV154** | leaf, root, sediment | none, none, above | *Alternaria alternata* |
| **ITS_SV260** | leaf, root | none, none | *Saccharomyces paradoxus* |
| **ITS_SV362** | leaf | below | *Lobulomyces* sp. |
| **ITS_SV381** | leaf | above | *Aureobasidium pullulans* |
| **ITS_SV389** | leaf | none | *Lecanora populicola* |
| **ITS_SV426** | leaf, sediment | none, none | *Saccharomyces* sp. |
| **ITS_SV779** | leaf, root, sediment | above, above, none | *Malassezia restricta* |

| | | | |
|---|---|---|---|
| **ITS_SV807** | leaf | none | *Meliniomyces* sp. |
| **ITS_SV1714** | leaf | above | Unclassified Capnodiales sp. |
| **ITS_SV2181** | leaf | none | Unclassified Capnodiales sp. |
| **ITS_SV2377** | leaf | none | Unclassified Capnodiales sp. |
| **ITS_SV5566** | leaf | none | *Malassezia globosa* |
| **ITS_SV60** | root | below | Unclassified Sordariomycetes sp. |
| **ITS_SV101** | root | none | Unclassified Didymellaceae sp. |
| **ITS_SV125** | root | none | Unclassified Ascomycota sp. |
| **ITS_SV169** | root | none | Unclassified Didymellaceae sp. |
| **ITS_SV234** | root | none | Unclassified Sordariomycetes sp. |
| **ITS_SV497** | root | below | Unclassified Sordariomycetes sp. |
| **ITS_SV540** | root | none | *Phaeotheca salicorniae* |
| **ITS_SV590** | root | above | *Cladosporium halotolerans* |
| **ITS_SV766** | root | none | *Phaeotheca salicorniae* |

| | | | |
|---|---|---|---|
| **ITS_SV1045** | root | above | *Hortaea werneckii* |
| **ITS_SV31** | sediment | above | *Cladosporium* sp. |
| **ITS_SV144** | sediment | none | *Cystobasidium pinicola* |
| **ITS_SV210** | sediment | above | *Didymella glomerata* |
| **ITS_SV404** | sediment | below | Unclassified Pleosporaceae sp. |
| **ITS_SV630** | sediment | above | *Paraphaeosphaeria angularis* |
| **ITS_SV679** | sediment | above | *Pseudeurotium bakeri* |
| **ITS_SV880** | sediment | below | Unclassified fungal sp. |
| **ITS_SV950** | sediment | above | *Pyrenochaetopsis leptospora* |
| **ITS_SV952** | sediment | none | Unclassified Sclerotiniaceae sp. |
| **ITS_SV962** | sediment | above | *Paraconiothyrium cyclothyrioides* |
| **ITS_SV1255** | sediment | above | *Wickerhamomyces anomalus* |
| **ITS_SV1468** | sediment | above | *Paraconiothyrium brasiliense* |
| **ITS_SV1538** | sediment | above | *Pyrenochaetopsis leptospora* |
| **ITS_SV1745** | sediment | above | *Trichoderma* sp. |

| | | | |
|---|---|---|---|
| **ITS_SV1928** | sediment | above | Unclassified Hyaloscyphaceae sp. |
| **ITS_SV2045** | sediment | above | *Saitozyma podzolica* |
| **ITS_SV2047** | sediment | above | *Psathyrella maculata* |
| **ITS_SV2164** | sediment | above | Unclassified Saccharomycetales sp. |
| **ITS_SV2245** | sediment | none | Unclassified Rhizophydiales sp. |
| **ITS_SV2430** | sediment | above | *Ganoderma* sp. |
| **ITS_SV2439** | sediment | above | *Penicillium* sp. |
| **ITS_SV2488** | sediment | none | *Trichoderma* sp. |
| **ITS_SV2529** | sediment | above | *Lycoperdon nigrescens* |
| **ITS_SV2601** | sediment | above | *Paraphaeosphaeria* sp. |
| **ITS_SV2610** | sediment | above | *Paraconiothyrium archidendri* |
| **ITS_SV2625** | sediment | above | *Paraconiothyrium* sp. |
| **ITS_SV2685** | sediment | above | Unclassified Pleosporales sp. |
| **ITS_SV2716** | sediment | none | *Lycoperdon pyriforme* |
| **ITS_SV2876** | sediment | none | Unclassified Aphelidiomycota |

| | | | |
|---|---|---|---|
| | | | sp. |
| **ITS_SV2885** | sediment | above | *Lachancea mirantina* |
| **ITS_SV3107** | sediment | above | Unclassified Teichosporaceae sp. |
| **ITS_SV3162** | sediment | above | *Didymocyrtis cladoniicola* |
| **ITS_SV3286** | sediment | above | *Penicillium* sp. |
| **ITS_SV3401** | sediment | above | *Coprinellus* sp. |
| **ITS_SV3483** | sediment | above | *Trichoderma* sp. |
| **ITS_SV3636** | sediment | above | *Fomitiporella* sp. |
| **ITS_SV3851** | sediment | above | Unclassified Microbotryales sp. |
| **ITS_SV4118** | sediment | none | *Hypholoma fasciculare* |
| **ITS_SV4193** | sediment | above | *Myrothecium* sp. |
| **ITS_SV5121** | sediment | above | *Trichoderma* sp. |
| **ITS_SV5439** | sediment | none | *Pyrenochaetopsis tabarestanensis* |
| **ITS_SV5656** | sediment | none | *Paraphaeosphaeria angularis* |
| **ITS_SV5741** | sediment | above | *Pyrenochaetopsis leptospora* |

| | | | |
|---|---|---|---|
| **ITS_SV5941** | sediment | none | *Cortinarius bivelus* |
| **ITS_SV6868** | sediment | above | *Endoxyla* sp. |
| **ITS_SV7233** | sediment | above | *Lophiostoma* sp. |
| **ITS_SV7652** | sediment | none | Unclassified Hypocreales sp. |
| **ITS_SV8621** | sediment | none | *Betamyces* sp. |
| **ITS_SV11477** | sediment | above | *Coprinellus micaceus* |
| **ITS_SV11928** | sediment | none | Chytridiomycetes sp. |
| **ITS_SV12885** | sediment | above | *Phialocephala humicola* |
| **ITS_SV13529** | sediment | above | *Phaeosphaeriaceae* sp. |
| **ITS_SV16671** | sediment | none | *Hypholoma capnoides* |
| **ITS_SV21975** | sediment | none | Unclassified Dictyosporiaceae sp. |
| **ITS_SV50231** | sediment | none | Unclassified Gigasporales sp. |

**Table S4.** Predicted core 18S rRNA gene ASVs. ASVs were ranked by their abundance-occupancy distribution and then predicted to be in the core based on a final percent increase of equal or greater than 10% to Bray-Curtis dissimilarity.The Sloan neutral model was then applied to the abundance-occupancy distributions to identify ASVs that deviate such that ASVs above the neutral model are predicted to be selected for by the environment (e.g. by the host plant, *Z. marina*), and those below the model are predicted to be selected-against or dispersal-limited. Here we report the core 18S rRNA gene ASVs for the *Z. marina* leaf, root and sediment mycobiomes, whether these ASVs deviate from the neutral model (above, below, none) and the taxonomy of each ASV.

| ASV | Core prediction | Neutral model deviation | Taxonomy |
|---|---|---|---|
| **18S_SV756** | leaf, root | none, none | Unclassified Chytridiomycetes sp. |
| **18S_SV928** | leaf, root | above, above | *Saccharomyces* sp. |
| **18S_SV968** | leaf | none | Unclassified Lobulomycetaceae sp. |
| **18S_SV1004** | leaf | none | *Saccharomyces* sp. |
| **18S_SV1553** | leaf | none | Unclassified Gromochytriaceae sp. |
| **18S_SV1567** | leaf, root | none, none | Unclassified Sordariomycetes sp. |
| **18S_SV1734** | leaf, root | none, none | Unclassified Rhizophydiales sp. |
| **18S_SV1977** | leaf, root | none, none | *Chytridium* sp. |

| | | | |
|---|---|---|---|
| **18S_SV3823** | leaf | none | *Cryptococcus* sp. |
| **18S_SV970** | root | none | Unclassified Sordariomycetes sp. |
| **18S_SV2546** | root | none | Unclassified Chytridiomycota sp. |
| **18S_SV3446** | root | none | Unclassified Aphelidea sp. |
| **18S_SV4479** | root | below | Unclassified Sordariomycetes sp. |
| **18S_SV5426** | root | above | Unclassified Chytridiomycota sp. |
| **18S_SV8081** | root | none | Unclassified Lobulomycetaceae sp. |
| **18S_SV10835** | root | none | *Chytridium* sp. |
| **18S_SV18125** | root | none | Unclassified Lobulomycetaceae sp. |
| **18S_SV27158** | root | none | Unclassified Chytridiomycetes sp. |
| **18S_SV897** | sediment | none | Unclassified Chytridiomycetes sp. |
| **18S_SV1751** | sediment | none | Unclassified Rhizophydiales sp. |
| **18S_SV2055** | sediment | above | *Trichoderma* sp. |
| **18S_SV2127** | sediment | none | Unclassified Spizellomycetales sp. |
| **18S_SV2138** | sediment | none | Unclassified Chytridiomycetes sp. |
| **18S_SV3230** | sediment | below | *Metschnikowia* sp. |

| | | | |
|---|---|---|---|
| **18S_SV5260** | sediment | none | *Metschnikowia* sp. |
| **18S_SV8806** | sediment | none | *Blastobotrys* sp. |
| **18S_SV14838** | sediment | below | Unclassified Boletales sp. |
| **18S_SV18689** | sediment | none | Unclassified Ascomycota sp. |
| **18S_SV20875** | sediment | none | Unclassified Rhizophydiales sp. |
| **18S_SV23021** | sediment | none | Unclassified Agaricales sp. |
| **18S_SV23280** | sediment | none | *Rhizophydium* sp. |
